# Profiling and Targeting of Regulatory RNAs to Upregulate Gene Expression

**DOI:** 10.64898/2026.04.21.719874

**Authors:** Brynn N. Akerberg, Bryan J. Matthews, Yuting Liu, Cecile Mathieu, Jenna Williams, Salome Manska, Jiaqi Huang, Yuichi Nishi, Abeer Almutairy, Eric Coughlin, Gabriel Golczer, Scott Waldron, Isabella Pellegrino, Yuchun Guo, Evan Cohick, Harpreet Turna, Kevin Xiong, Gavin Whissell, Rachana S. Kelkar, Mario Gamboa, Evan Lenz, Christopher J. Jurcisin, Rutuja Pai, Justin A. Caravella, Alfica Sehgal, Daniel F. Tardiff, Alla A. Sigova, David Bumcrot

**Affiliations:** CAMP4 Therapeutics Corp., Cambridge, MA USA

## Abstract

Transcription of long noncoding RNAs (lncRNAs), including enhancer RNAs (eRNAs) and promoter-associated RNAs (paRNAs), collectively termed *regulatory RNAs* (regRNAs), is a hallmark of active gene expression, yet it remains unknown whether regRNAs can be targeted to selectively enhance transcription *in cis*. We developed regRNA Capture-seq, a high-throughput method to profile regRNAs, and applied it to primary human hepatocytes, annotating thousands of regRNAs at ∼2,000 enhancers and promoters. Using this approach, we interrogated a genetically validated enhancer of the *ornithine transcarbamylase* (*OTC*) gene, mutations of which cause OTC deficiency (OTCD), the most common urea cycle disorder. Antisense oligonucleotides (ASOs) targeting enhancer-derived regRNAs led to dose-dependent upregulation of *OTC* in hepatocytes. Mechanistically, ASOs altered regRNA structure, elevated regRNA levels, displaced transcriptional repressors, and increased H3K27 acetylation at the targeted enhancer.

This work establishes a potential therapeutic strategy for addressing haploinsufficiency and highlights regRNAs as actionable targets for ASO-mediated upregulation of gene expression.

## Introduction

Transcription factors (TFs) regulate gene expression by binding to specific DNA motifs within proximal and distal regulatory elements, including promoters and enhancers. These interactions facilitate the spatial organization of chromatin, bringing regulatory elements into close proximity and promoting the recruitment of the RNA polymerase II (RNA Pol II) complex. At promoters, RNA Pol II initiates bidirectional transcription from transcription start sites (TSSs), generating both protein-coding messenger RNAs (mRNAs) and non-coding promoter-associated RNAs (paRNAs). In contrast, transcription at enhancers primarily produces non-coding enhancer RNAs (eRNAs)^1,2^.

Cell-type-specific gene expression programs are largely orchestrated by the selective activation of tens of thousands of enhancers^3^. These regulatory elements govern the production of diverse mRNA species and hundreds of thousands of non-coding RNAs, shaping the unique transcriptomes that define cell identity and function. The dynamic regulation of gene expression through enhancer activity underlies key biological processes, including development, responses to external stimuli, and tissue homeostasis. Individual genes can be regulated by multiple enhancers, many of which act with high specificity and exert minimal influence on other loci^4–7^.

Emerging evidence suggests that, beyond their canonical DNA-binding roles, TFs and cofactors also engage in functional interactions with nascent non-coding RNAs, such as eRNAs and paRNAs, collectively termed *regulatory RNAs* (regRNAs), at active regulatory elements^1,8–10^. These RNA-protein interactions can locally concentrate TFs and cofactors near their binding sites, enhancing the probability of TF rebinding and influencing both transcription initiation and elongation. Through these mechanisms, regRNAs can modulate enhancer and promoter activity, fine-tune gene expression, and contribute to the robustness and plasticity of transcriptional programs.

Most eRNAs and paRNAs are unspliced and non-polyadenylated, reflecting the absence of canonical splicing and polyadenylation signals at their genomic loci^11–13^. Consequently, these transcripts are highly unstable, with half-lives of only a few minutes^13–15^, and are rarely detected in steady-state RNA pools. Even when spliced and polyadenylated, regRNAs remain significantly less stable than mRNAs^16^, posing substantial challenges for their reliable detection, quantification, and functional characterization.

Due to their low abundance and inherent instability, regRNAs have remained challenging to annotate and functionally characterize^17^. In contrast to long intergenic non-coding RNAs (lincRNAs), which are typically more stable and readily detectable^18^, eRNAs and paRNAs require specialized detection strategies. Techniques like Precision Run-On sequencing (PRO-seq) and Global Run-On sequencing (GRO-seq) enable the identification of nascent, unstable transcripts by capturing RNA associated with actively elongating RNA Pol II^19^. These approaches have provided key insights into transcriptional dynamics at regulatory regions. However, they do not capture full-length transcript information, limiting their utility for studying regRNA structure and downstream function.

Conventional RNA sequencing methods, including those incorporating ribosomal RNA depletion, also lack the sensitivity to reliably detect these unstable transcripts^20^. As a result, improved strategies are needed to enhance the annotation, detection, and mechanistic interrogation of regRNAs to better understand their roles in transcriptional regulation and cell-type-specific gene expression programs.

Substantial progress has been made in developing loss-of-function strategies to interrogate the functional roles of long non-coding RNAs (lncRNAs), including lincRNAs, paRNAs, and eRNAs, at both the transcriptional and post-transcriptional levels. These approaches include the use of transcriptional inhibitors, small interfering RNAs (siRNAs), antisense oligonucleotides (ASOs), and the insertion of premature polyadenylation signals to terminate transcription^21–23^. In many cases, perturbation of these non-coding RNAs leads to a reduction in expression of their associated target genes, suggesting a positive regulatory relationship between regRNA production and gene activation. Further evidence for this correlation comes from studies showing that transcriptional induction of eRNAs frequently precedes the upregulation of nearby protein-coding genes in response to external stimuli^24^. Moreover, genes associated with highly transcriptionally active enhancers, particularly those classified as super-enhancers, tend to exhibit elevated expression levels, reinforcing the idea that regRNAs contribute to enhancer-mediated gene activation^25–29^.

Many loss-of-function or hypomorphic mutations in protein-coding genes result in haploinsufficiency and associated diseases. In such cases, even modest increases in expression, often less than two-fold of the remaining functional allele, could be therapeutically beneficial^30^. This raises the possibility that enhancing the expression of disease-associated genes could be achieved through the targeted induction of regRNAs. However, previous attempts to deliver exogenous regRNAs *in trans* have produced inconsistent results, likely due to the predominantly *cis*-acting nature of these transcripts, which function at or near their site of transcription^31^. These limitations highlight the need for new strategies to selectively induce regRNA expression at endogenous loci, with the goal of modulating enhancer activity and therapeutically boosting target gene expression.

Here, we present a novel high-throughput regRNA Capture-seq method for annotating eRNAs and paRNAs in primary human hepatocytes (PHHs). We applied this approach to characterize regRNAs transcribed from a genetically validated enhancer of the *ornithine transcarbamylase* (*OTC*) gene, the most frequently mutated gene in urea cycle disorders (UCDs). A targeted ASO screen against enhancer-derived regRNAs identified multiple ASOs capable of inducing concentration-dependent upregulation of *OTC* expression in PHHs. Mechanistic studies revealed that ASO binding induces structural remodeling of the regRNA, leading to eviction of negative transcriptional regulators from the enhancer and increased transcriptional activity. These findings establish a proof-of-concept for therapeutic gene activation through targeted modulation of regRNAs.

## Results

### regRNA catalog in primary human hepatocytes

We have compiled a catalog of polyadenylated and non-polyadenylated regRNA species transcribed from thousands of enhancers in PHHs, as shown in Figure 1A. To capture these low-abundance non-coding RNAs, we developed a novel method, regRNA Capture-seq, that combines target capture with long-read sequencing. This approach uses biotinylated RNA probes to selectively hybridize with regRNAs through Watson-Crick base pairing, allowing their pulldown from total RNA isolated from PHHs using streptavidin beads. A similar technique has been previously employed for targeted enrichment of specific genomic DNA loci, as well as mRNA, rRNA, snRNA, and lincRNAs, which are more abundant than regRNAs^32–34^. The key advantages of our method include its high sensitivity, versatility, and throughput, enabling the detection of non-coding RNAs transcribed from individual enhancers or promoters, or even from hundreds or thousands of enhancers and promoters simultaneously.

**Figure 1.**
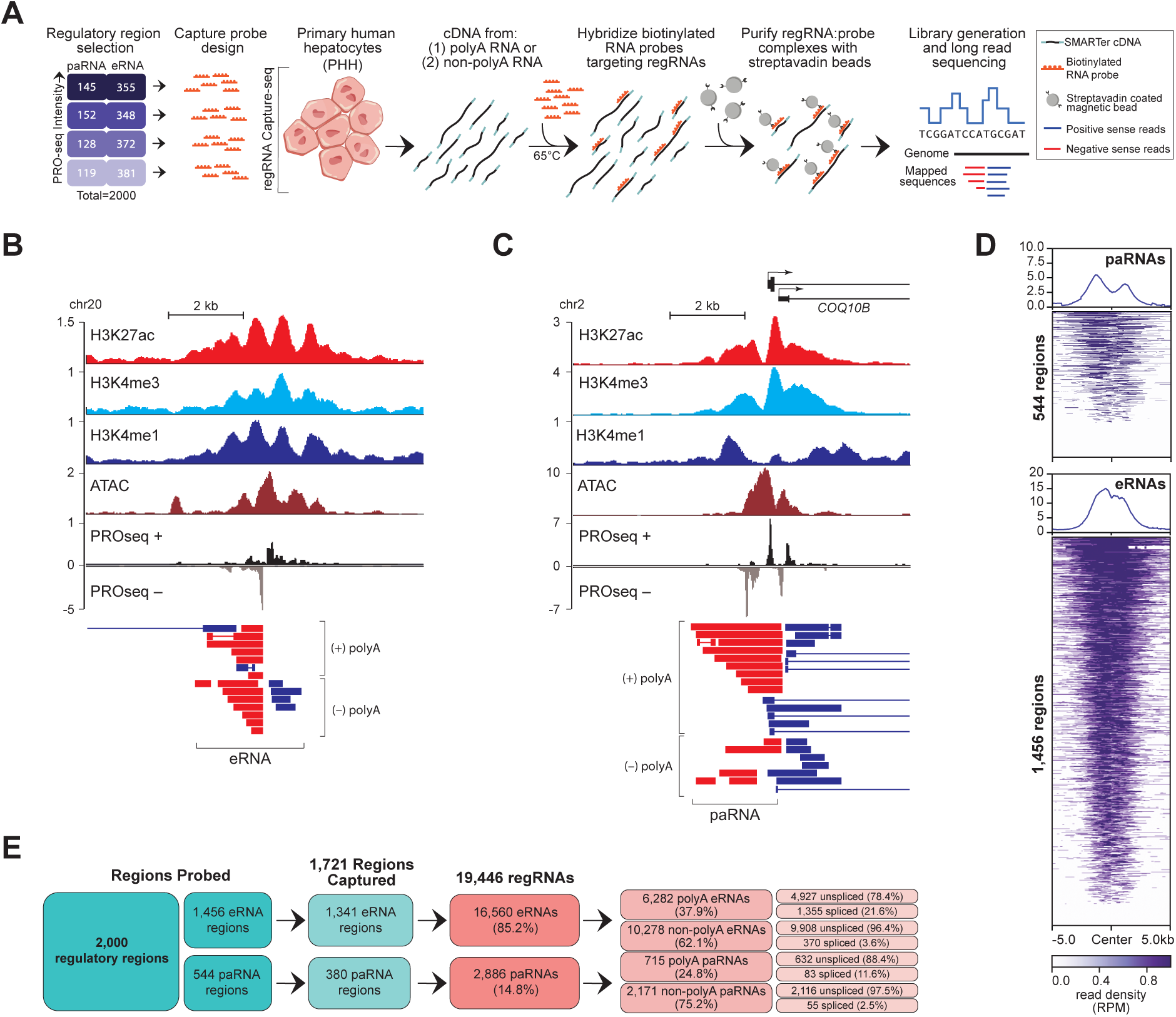
Regulatory RNA catalog in primary human hepatocytes. (A) Schematic of the strategy used to generate a catalog of polyadenylated and non-polyadenylated regRNA species transcribed from ∼2,000 targeted enhancers in PHHs. (B) Representative example of regRNA species transcribed from an intergenic enhancer located ∼40 kb upstream of the *PTPN1* gene. Signal tracks show regRNA Capture-seq species alongside ChIP-seq for H3K27Ac, H3K4me3, and H3K4me1, as well as ATAC-seq and PRO-seq signals. Signal intensity is shown in reads per million (RPM). regRNAs mapping to the plus strand are shown in blue, minus strand in red. PRO-seq reads mapping to the plus strand are shown above the x-axis, minus strand below. (C) Example of regRNA species detected at the promoter of the *COQ10B* gene. regRNA reads (red) and mRNA reads (blue) are shown with corresponding chromatin and nascent transcription profiles as in (B). Signal intensity is shown in RPM. PRO-seq reads mapping to the plus strand are shown above the x-axis, minus strand below. (D) Heatmaps showing the distribution of regRNA reads ±5 kilobases (kb) from ATAC-seq peak centers at 1,456 eRNA-producing regions (bottom) and 544 paRNA-producing regions (top). Reads mapping to Watson and Crick strands are combined and shown together in purple. mRNA reads are excluded. Color bar scale represents normalized read counts per 10 nt bin. (E) Summary flowchart illustrating the classification and key features of regRNA species identified by regRNA Capture-seq in PHHs.

To create a comprehensive catalog of regRNAs in PHHs, we integrated Assay for Transposase-Accessible Chromatin with sequencing (ATAC-seq) data with Chromatin Immunoprecipitation sequencing (ChIP-seq) data for histone 3 acetylated at lysine 27 (H3K27Ac). This analysis identified 134,520 active enhancers, defined as ATAC-seq peak summits overlapping H3K27Ac-enriched regions within ± 500 nucleotides (nt) (Figure S1A). Notably, 68% of these enhancers were intragenic (located within the body of protein-coding genes), while the remaining ones were located outside annotated gene boundaries (Figure S1B). Closer examination of the intragenic subset revealed that 14.2% were associated with gene promoters (within ± 1000 nt of TSS), suggesting potential regulatory interplay between enhancer and promoter activity (Figure S1B). To assess the transcriptional output of these enhancers, we performed PRO-seq, which detects nascent RNA associated with actively elongating RNA Pol II. Beyond mapping the positions of engaged RNA Pol II, PRO-seq provides strand-specific quantification of nascent transcripts, offering insights into both the orientation and relative abundance of regRNA transcription. Unlike conventional RNA-seq, PRO-seq uniquely captures transient regRNAs, which are protected from degradation during RNA Pol II elongation. Consistent with observations in other cell types, the majority of active enhancers (96,169 of 134,520) displayed detectable transcriptional activity, as defined by PRO-seq signal exceeding a threshold of 0.02 reads per million (RPM) (Figure S1C). We next compared PRO-seq read densities at enhancers (not overlapping promoters) with PRO-seq signals upstream of gene promoters (Figures S1D and S1E). Analysis of the top 10,000 eRNA– and paRNA-producing regions, ranked by read density, revealed comparable transcriptional activity at both (Figure S1D), suggesting similar levels of regRNA output. Notably, the average PRO-seq read density for both eRNAs and paRNAs was higher than the median, indicating a skewed distribution, in which a small subset of highly active regulatory elements contributes disproportionately to the overall transcriptional output (Figure S1E).

To characterize regRNAs transcribed from regulatory regions with varying transcriptional activities, as defined by PRO-seq, we stratified hepatocyte enhancers into four subgroups: very highly active, highly active, moderately active, and lowly active (Figure S2A). Each subgroup was profiled independently to avoid signal saturation and minimize overrepresentation of highly abundant regRNAs relative to less abundant species. An equal number of enhancers were selected from each subgroup for the design of biotinylated RNA probes (Figure 1A). In total, ∼125,000 capture probes were designed: four sets of ∼31,000 probes corresponding to each of the four enhancer subgroups to enrich for regRNAs produced by ∼2,000 enhancers with differing expression levels (Figure S2A). These enhancers encompassed both intra– and intergenic regions and included loci producing eRNAs and paRNAs, enabling a comprehensive analysis that was not biased toward any specific class of regRNAs (Figure S2B).

Total RNA was isolated from PHHs and separated into polyadenylated and non-polyadenylated fractions. Both fractions were converted into cDNA, amplified, and hybridized separately with each of the four sets of capture probes. The isolated cDNA was then further amplified and sequenced using long-read sequencing, enabling the identification of full-length polyadenylated and non-polyadenylated regRNAs (Figure 1A). This approach preserved detailed information about the regRNA repertoire at individual enhancers (Figures 1B and 1C). Eighty-six percent (1,721) of enhancers had detectable regRNAs, indicating that regRNA Capture-seq achieved sensitivity comparable to PRO-seq (Figures 1D, 1E, and S2C). Despite their comparable sensitivity, PRO-seq and regRNA Capture-seq exhibited markedly different signal distributions at enhancers (Figures 1D and S2D), suggesting they capture complementary, rather than redundant, aspects of regRNA transcription. Both methods revealed peaks of bidirectional transcription in aggregate plots of the selected group of ∼2,000 enhancers. However, PRO-seq signals were narrower and concentrated within ±500 nt of the enhancer center, consistent with the detection of nascent regRNA transcripts associated with paused RNA Pol II. In contrast, regRNA Capture-seq signals were significantly broader, extending up to ±2,000 nt from the enhancer center, indicative of full-length regRNA transcript detection (Figures 1D and S2D). The overall regRNA coverage included 19,446 distinct transcripts, with a higher representation of non-polyadenylated species (12,449) compared to polyadenylated ones (6,997) (Figure 1E). Importantly, the majority of evaluated enhancers (58.8%, 1,176) produced both polyadenylated and non-polyadenylated regRNAs, with an average of seven regRNAs per enhancer (Figure S2E). Consistent with the stratified experimental design, the total number of detected eRNA and paRNA species did not differ substantially between highly transcribed enhancers (very high and high) and those with moderate or low activity (Figure S2F). However, the number of long reads supporting individual eRNA and paRNA annotations was higher in the very high and high activity groups compared to the moderate and low groups (Figure S2G), suggesting that regRNA Capture-seq can detect relative differences in regRNA abundance. These differences, however, were semi-quantitative and less precisely resolved than those measured by PRO-seq (Figures S2A and S2G).

In contrast to previous reports on lincRNAs, only 32.7% of the regRNA species detected in PHHs were multi-exonic, and just 29.3% of these contained canonical splicing motifs. These data suggest that only ∼9.6% of the regRNAs may undergo conventional splicing (Data Table 1). Notably, only ∼32% of unspliced regRNAs were found in the polyadenylated fraction, while ∼68% of unspliced regRNAs were found in the non-polyadenylated fraction, suggesting that unspliced regRNAs are typically not polyadenylated (Figure 1E). As expected, polyadenylated regRNAs were generally longer than their non-polyadenylated counterparts (median of 1,246 nt vs. 906 nt) (Figure S2H). However, greater heterogeneity was observed at the 3’ end of regRNAs, which contrasts with the 5’ heterogeneity typically seen in mRNAs that are degraded in the 5’ to 3’ direction (Figure S2I, S2J). Even unspliced polyadenylated regRNAs were longer than non-polyadenylated species (median of 1,101 nt vs. 866 nt) (Figure S2K), suggesting that the detected non-polyadenylated transcripts were likely either nascent regRNAs or degradation products of longer regRNA species, which are expected to be degraded in the 3’ to 5’ direction by the nuclear exosome complex. In contrast to earlier reports, we did not detect short ∼100 nt eRNA species, which are typically premature termination products of paused RNA Pol II detected by other methods^35^. This may be due to their rapid degradation compared to other non-polyadenylated RNA species.

Closer inspection of regRNAs at individual enhancers revealed that 45.6% (785 of 1,721) exhibited bidirectional transcription, while 54.4% (936 of 1,721) showed unidirectional transcription, with regRNAs mapping exclusively to either the Watson (plus) or Crick (minus) strand of DNA. As expected, approximately 40.6% (380 of 936) of unidirectional transcripts were paRNAs originating upstream of gene promoters. Further analysis of regRNA Capture-seq and PRO-seq signals at the remaining 59.4% (556 of 936) of unidirectional enhancers showed significantly fewer reads compared to bidirectional enhancers (Figure S2L), indicating that unidirectional enhancers are generally less transcriptionally active than their bidirectional counterparts.

Remarkably, in addition to correctly identifying 642 previously annotated lincRNAs, regRNA Capture-seq identified 19,446 novel regRNA species, representing 96.8% of the PHH catalog, that have not been previously annotated (Data Table 2).

### *OTC* enhancer is a transcriptionally active regulatory element in cultured hepatocytes and liver tissue

To evaluate the functional role of regRNAs and their potential as therapeutic targets, we focused on a previously identified regulatory region upstream of *ornithine transcarbamylase* (*OTC*), a key gene in the urea cycle. Mutations in any of the six enzymes involved in the urea cycle can contribute to UCDs. However, the majority of UCD patients carry mutations in the coding regions of *OTC*, which reduce its enzymatic activity, leading to *OTC* deficiency (OTCD). This condition is characterized by the systemic accumulation of toxic ammonia and impaired urea production^36^.

Recent studies have demonstrated that a subset of OTCD patients harbor mutations in the liver-specific enhancer of *OTC*, located approximately 9 kilobases (kb) upstream of the *OTC* promoter^37^. This enhancer is evolutionarily conserved across mammalian species, and mutations within this region lead to reduced *OTC* expression. These findings strongly suggest that such mutations could contribute to the development of OTCD.

To further characterize the upstream *OTC* enhancer, we analyzed ATAC-seq, H3K27Ac, and H3K4me3 ChIP-seq signals at the enhancer and promoter regions in PHHs and human liver tissue (Figure 2A, top). While both H3K27Ac and H3K4me3 are associated with active regulatory regions, H3K4me3 is more commonly enriched at promoters, whereas H3K27Ac marks active promoters and enhancers^38^. Our analysis confirmed the presence of an active *OTC* enhancer in both PHHs and human liver tissue. Consistent with previous reports that cooperative binding of hepatocyte nuclear factor 4 A (HNF4A) and CCAAT/enhancer-binding protein beta (C/EBPβ) confers liver-specific activity to the *OTC* enhancer^37,39^, we identified binding motifs for both factors within the enhancer region (Figure S3A). In addition, we detected motifs for other hepatocyte-enriched TFs, including FOXA1 and FOXA2, as well as for TFs expressed in PHHs and human liver, such as signal transducer and activator of transcription 1 (STAT1) (Figure S3A). These findings suggest that these TFs may contribute to *OTC* enhancer function and regulation of *OTC* gene expression.

**Figure 2.**
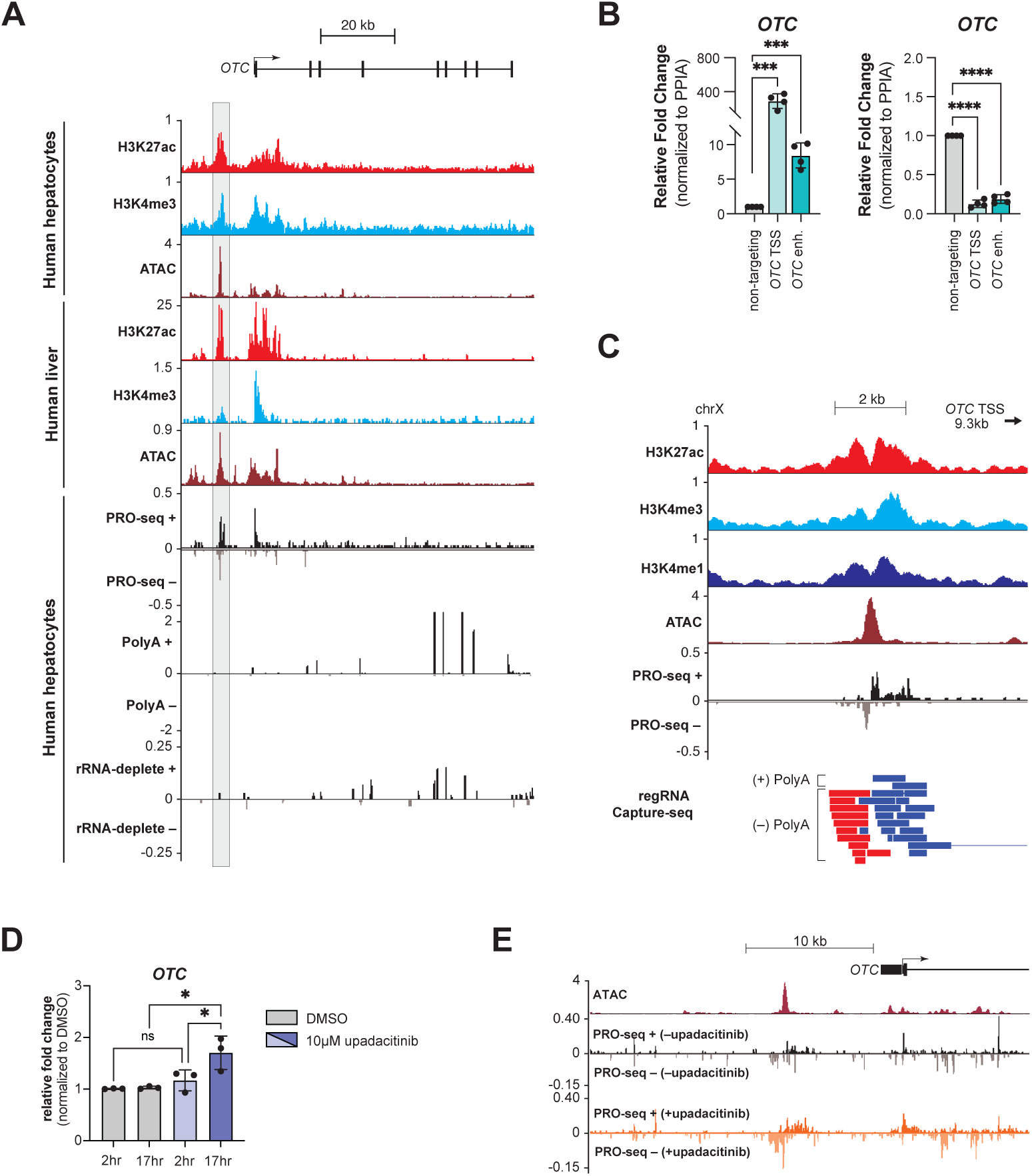
The *OTC* enhancer is a transcriptionally active regulatory element in cultured hepatocytes and liver tissue. (A) Signal tracks at the *OTC* locus in PHHs and human liver tissue, showing ChIP-seq profiles for H3K27Ac and H3K4me3, along with ATAC-seq, PRO-seq, and RNA-seq from polyA-selected and ribo-depleted total RNA. PRO-seq and RNA-seq reads mapping to the plus strand are shown above the x-axis, minus strand below. *OTC* enhancer is highlighted with a gray box. Signal intensity is shown in RPM. (B) qRT-PCR analysis of *OTC* mRNA expression following CRISPR activation (CRISPRa) and interference (CRISPRi) using dCas9-VPH (left) or dCas9-KRAB (right), correspondingly, targeted to the *OTC* TSS or enhancer in HepG2 cells. A non-targeting sgRNA served as a negative control. Data are presented as mean fold change for *OTC* relative to peptidylprolyl isomerase A (*PPIA*) ± s.d. (n = 4). Statistical significance was assessed by unpaired student’s *t*-test (***, p<0.0005; ****, p<0.0001). (C) regRNA species transcribed from the *OTC* enhancer in PHHs. Signal tracks show regRNA Capture-seq reads along with ChIP-seq for H3K27Ac, H3K4me3, and H3K4me1, and corresponding ATAC-seq and PRO-seq profiles. Signal intensity is shown in RPM. regRNA reads mapping to the plus strand are shown in blue, minus strand in red. PRO-seq reads mapping to the plus strand are shown above the x-axis, minus strand below. (D) *OTC* mRNA expression measured by qRT-PCR following 2 h and 17 h treatment of PHHs with 10 µM upadacitinib. DMSO served as a control. Data represent mean fold change relative to *PPIA* ± s.d. (n = 3). Statistical significance was determined by one-way ANOVA (*, p<0.01 and 0.04 upadacitinib at 2 h v. 17 h and DMSO v. upadacitinib at 17 h, respectively). (E) PRO-seq signal at the *OTC* locus in PHHs treated with DMSO or 10 µM upadacitinib, shown alongside ATAC-seq signal. PRO-seq reads from the plus strand are shown above the x-axis, minus strand below. Signal intensity is shown in RPM.

Using PRO-seq data, we observed a classical pattern of divergent transcription at the *OTC* enhancer in PHHs, with similar signal intensity upstream and downstream of the enhancer center, which contains predicted binding sites for multiple liver-specific TFs (Figure 2A, bottom). However, no signal was detected at the enhancer in the polyA-selected or ribodepleted RNA-seq datasets from PHHs. These findings indicate that the *OTC* enhancer is transcriptionally active in cultured hepatocytes, although the resulting eRNAs do not accumulate to levels detectable by conventional RNA-seq.

To directly evaluate the functional contribution of the *OTC* enhancer to gene expression, we used guide RNAs to target the enhancer region in HepG2 cells, where the *OTC* gene is expressed and the enhancer is functional (Figure S3B). We employed a catalytically inactive Cas9 (dCas9) fused to either the VPH activation^40^ or KRAB repression domain^41^. As expected, targeting the enhancer or promoter with these constructs led to corresponding increases or decreases in *OTC* mRNA levels, respectively, with the largest effect size observed at promoters, as previously reported^40,42^. In total, this supports a direct regulatory role for the enhancer in controlling *OTC* gene expression (Figure 2B).

To characterize the regRNAs produced from the enhancer, we performed regRNA Capture-seq in PHHs and human liver tissue using capture probes designed specifically for the *OTC* enhancer (Figure S3D). This approach revealed a diverse set of predominantly unspliced polyadenylated and non-polyadenylated transcripts originating from both the Watson (plus) and Crick (minus) strands of the enhancer, consistent with divergent transcription (Figure 2C, Supplementary Table 7).

Notably, polyadenylated regRNA species were more abundant and diverse on the plus strand, whereas minus-strand regRNAs were primarily non-polyadenylated. This strand bias was supported by the polyadenylation signal analysis: plus-strand transcripts terminated 10–60 nt downstream of canonical polyadenylation motifs, while the nearest such motif on the minus strand was located over 350 nt away. Polyadenylated regRNAs had an average length of ∼880 nt, whereas non-polyadenylated transcripts were shorter, averaging ∼700 nt. Coding potential analysis confirmed that these transcripts are non-coding, with scores characteristic of lncRNAs^43^ (Figure S3C).

Unexpectedly, we also identified longer, spliced polyadenylated transcripts initiating from the enhancer and extending into the *OTC* gene body. These isoforms closely resembled the canonical *OTC* mRNA but included one or two upstream non-coding exons that are absent from the annotated transcript (Figure S3E). The extended transcripts contained canonical splice junctions, and qRT-PCR using exon junction-specific primers confirmed their presence in both PHHs and human liver tissue (Figure S3F). These results suggest that the *OTC* enhancer may also function as an alternative promoter, producing extended *OTC* transcripts that initiate at the same TSS as the plus-strand regRNAs and overlap with them over the first ∼1,600 nt.

To investigate the functional relationship between *OTC* enhancer-derived regRNAs and *OTC* mRNA expression, we pharmacologically modulated enhancer activity. Signaling pathway perturbation offers a broadly applicable strategy to evaluate the relationship between regRNA dynamics and transcriptional output, as reflected by coordinated changes in eRNA and target gene expression^27^. Given the central role of the AMP-activated protein kinase (AMPK) pathway in liver physiology^44^ and regulation of *OTC* gene expression^45^, we activated this pathway in PHHs using upadacitinib. In addition to being an inhibitor of the Janus kinase-signal transducer and activator of transcription (JAK-STAT) pathway, upadacitinib selectively promotes the expression and phosphorylation of AMPK^46^, leading to upregulation of *OTC* in hepatocytes^45^. Treatment of PHHs with upadacitinib resulted in differential expression of numerous genes, including upregulation of *OTC* (Figures 2D and S3G). Concurrently, PRO-seq analysis revealed increased nascent transcription at both the *OTC* enhancer and promoter regions (Figure 2E). These findings reveal a positive correlation between *OTC* regRNA production and gene transcription, consistent with a model in which *OTC* regRNAs directly promote *OTC* expression.

Together, these findings establish the *OTC* enhancer as a transcriptionally active regulatory element in both PHHs and liver tissue. It produces bidirectional non-coding RNAs that are positively associated with *OTC* mRNA levels and may also function as an alternative promoter, underscoring its central role in the transcriptional regulation of *OTC*.

### Identification and optimization of antisense oligonucleotides upregulating *OTC* gene expression in primary hepatocytes

Given the functional significance of the *OTC* enhancer in regulating *OTC* gene expression, we hypothesized that targeted upregulation of *OTC* enhancer-derived regRNAs at their site of transcription could increase *OTC* transcript levels. As most *OTC* mutations retain partial enzymatic activity, even modest increases in expression could restore enzyme function to the healthy range, potentially offering a therapeutic strategy for UCDs.

Oligonucleotides have become versatile tools for both functional genomics and therapeutic targeting of nucleic acid sequences. Among these, double-stranded small interfering RNAs (siRNAs) mediate gene silencing via sequence-specific hybridization, recruitment of the RNA-induced silencing complex (RISC), and subsequent cleavage of the target mRNA. In contrast, single-stranded antisense oligonucleotides (ASOs) bind to mRNA and recruit RNase H1, which recognizes RNA-DNA hybrids and catalyzes transcript degradation. ASOs are active in both the cytoplasm and nucleus, enabling them to function across cellular compartments where RNase H1 is present^47^. Moreover, certain chemically-modified ASOs act as steric blockers, interfering with RNA-binding proteins or spliceosomal components without inducing degradation^48,49^. These steric effects can alter transcript stability, splicing, or, in some contexts, promote transcript accumulation^50,51^.

Despite the growing use of ASOs in mRNA regulation, their application to regRNAs, particularly those transcribed from enhancers, remains largely unexplored. To date, only a few studies have demonstrated ASO-mediated downregulation of eRNAs^25,27,52^, leaving their broader therapeutic potential untapped.

To explore the feasibility for ASO-based modulation of *OTC* enhancer-derived regRNAs, we designed a panel of single-stranded, 20-nt-long ASOs complementary to regRNA sequences transcribed from both the Watson (plus) and Crick (minus) strands of the *OTC* enhancer that were identified by the regRNA Capture-seq in PHHs (Figure 3A, Supplementary Table 3). When designing the ASOs, we made no mechanistic assumptions about how ASOs might modulate the *OTC* regRNAs to allow for an unbiased assessment. As a result, this panel included both gapmer ASOs, incorporating chemical modifications that could support RNase H1-mediated cleavage, and fully chemically-modified ASOs that are thought to act through alternative mechanisms, including steric hindrance (Figure S4A). Non-targeting control ASOs with matched chemistries and minimal complementarity to *OTC* regRNAs were included for normalization. ASOs were delivered to PHHs via gymnotic uptake, a method that avoids transfection reagents and better mimics *in vivo* delivery. Following gymnotic delivery into PHHs at a concentration previously established to be effective for mRNA-targeting ASOs^51,53^ (5 µM), the majority of ASOs either had no measurable effect, or induced < 1.6-fold upregulation of *OTC* mRNA, regardless of the targeted strand or ASO chemistry (Figures 3B and S4A). However, a distinct subset of seven ASOs produced > 1.6-fold *OTC* mRNA upregulation. Further analysis revealed that this effect was reproducible, concentration-dependent, and specific for five of these ASOs, as none altered expression of the neighboring gene *RPGR* (Figures 3C and S4B).

**Figure 3.**
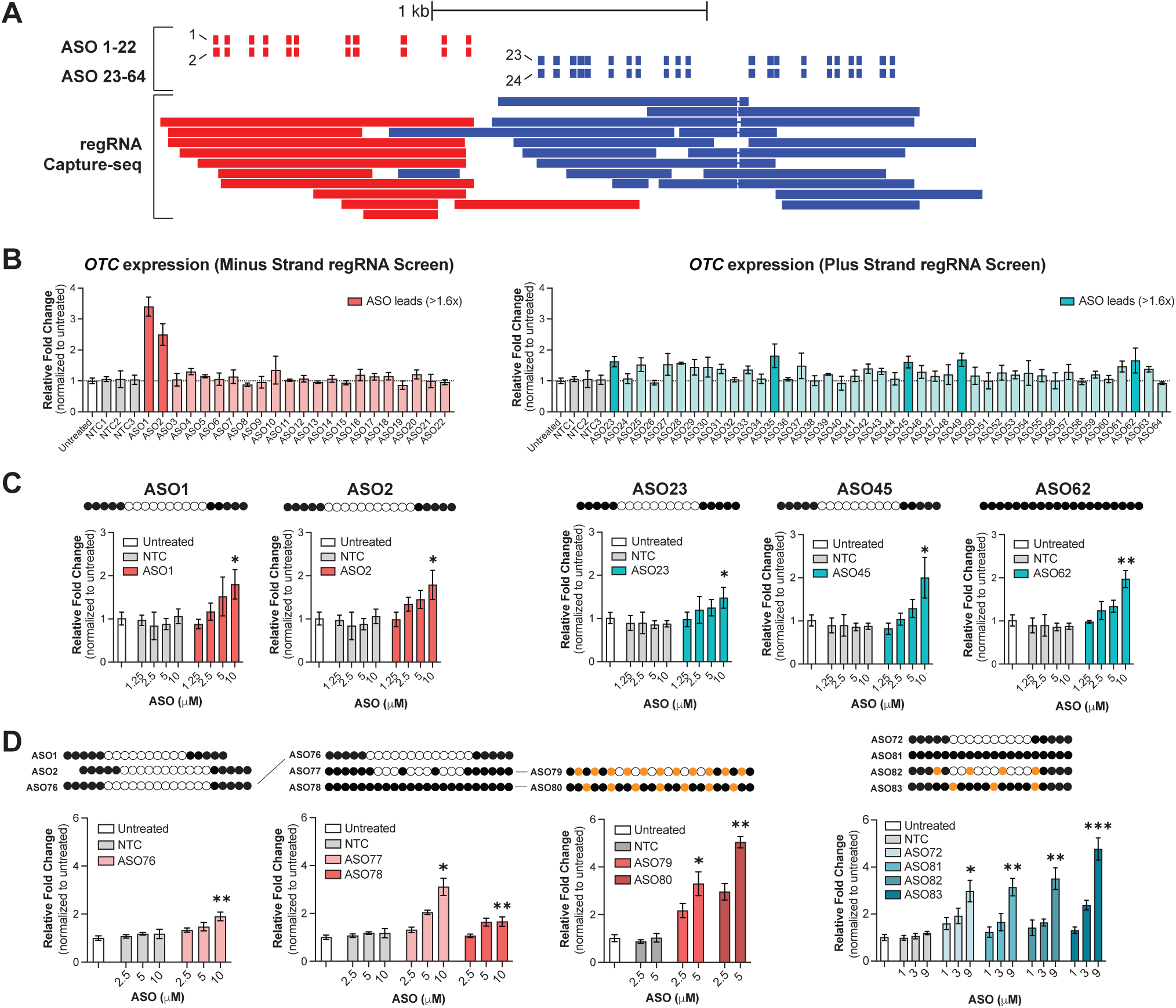
Identification and optimization of antisense oligonucleotides upregulating *OTC* expression in primary hepatocytes. (A) Schematic showing the positions of ASOs used in the screen, targeting *OTC* regRNAs transcribed from the minus (red) and plus (blue) strands of the *OTC* enhancer. (B) *OTC* mRNA expression measured by qRT-PCR following treatment of PHHs with 5 µM ASOs targeting either minus-strand regRNAs (left) or plus-strand regRNAs (right). Untreated cells and PHHs treated with three different sequence– and chemistry-matched non-targeting control (NTC) ASOs were used as controls. Data represent mean fold change relative to beta-2-microglobulin (*B2M)* ± s.d. (n = 3). (C) Dose-response curves of *OTC* mRNA expression following treatment of PHHs with increasing concentrations of ASOs targeting minus-strand (left) or plus-strand (right) regRNAs. NTC ASO-treated cells were used as controls. Data are shown as mean fold change for *OTC* relative to *B2M* ± s.d. (n = 3). ASO designs are illustrated above the graphs. Black circles represent 2′-MOE-modified nucleotides, white circles denote DNA nucleotides, and orange circles indicate LNA-modified nucleotides. Statistical significance was assessed by one-way ANOVA (*, p<0.05). (D) Dose-response curves of *OTC* mRNA expression following treatment with sequence– and chemistry-optimized ASOs targeting minus-strand (left) or plus-strand (right) regRNAs. NTC ASO-treated cells were used as controls. Data represent mean fold change relative to *B2M* ± s.d. (n = 3). ASO designs are illustrated above the graphs. Chemical modifications are represented as in Figure 3C.

These findings reveal a distinct, sequence-dependent, and potentially therapeutically relevant mechanism of action for this subset of ASOs. Notably, the five most active compounds included both steric-blocking and gapmer designs (Figure 3C), indicating that *OTC* mRNA upregulation is not restricted to a specific ASO chemistry. Instead, these data suggest that ASO effects might not require RNase H1-mediated cleavage and may be compatible with different types of ASO designs, underscoring the critical interplay between ASO sequence and chemistry in achieving effective regulatory outcomes. Importantly, two of the top-performing ASOs targeted regRNAs transcribed from the minus strand of the *OTC* enhancer and three targeted plus-strand regRNAs, demonstrating that regRNAs arising from either strand can control *OTC* expression.

To further enhance the efficacy of top-performing ASOs, we explored the design space through subtle sequence modifications to fine-tune binding site positioning, coupled with targeted adjustments to chemical modification patterns. Due to the overlap between sequences of ASO1 and ASO2 (Figure 3A), a single 23-mer ASO (ASO76) that included the sequences of both ASOs was designed and tested in PHHs. The ASO retained activity comparable to both parent compounds (Figure 3D, left). We observed a cluster of 3 gapmers inducing >1.5-fold *OTC* upregulation near the 5’ end of the plus-strand regRNA, tested additional ASOs in the region, and identified ASO72 with improved efficacy (Figures 3D, right and S4C). For gapmers targeting either the minus– or plus-strand regRNAs, conversion to partially modified mixmer designs (ASOs 77, 79, and 82) or fully modified steric-blocking chemistries (ASOs 78, 80, 81, and 83) further enhanced potency and efficacy (Figure 3D). Moreover, fully modified steric-blocking ASOs incorporating a combination of 2′-O-methoxyethyl (2′-MOE) and Locked Nucleic Acid (LNA) modifications (ASOs 80 and 83) demonstrated the greatest upregulation of *OTC* mRNA (>4-fold) (Figure 3D), potentially reflecting enhanced target affinity or improved gymnotic uptake. These findings further support the notion that RNase H1-dependent cleavage may not be required for ASO-dependent *OTC* upregulation. The effects of these ASOs were reproducible and concentration-dependent (Figure 3D).

To assess whether ASO-mediated *OTC* mRNA upregulation is consistent across multiple hepatocyte donors, we selected two top-performing ASOs, ASO80 and ASO83, targeting regRNAs on the minus and plus strands, respectively. Following gymnotic delivery, both ASOs consistently induced *OTC* mRNA expression across multiple PHH donors (Figure S4D), confirming that the stimulatory effects are donor-independent.

Together, these findings demonstrate that delivery of regRNA-targeting ASOs to PHHs can elicit sequence-specific upregulation of *OTC* expression. Furthermore, ASO efficacy can be further enhanced through empirically guided optimization of both sequence and chemical modifications.

### Mechanistic characterization of ASOs that upregulate *OTC* expression in PHHs

To investigate the mechanisms underlying the effects of regRNA-targeting ASOs on *OTC* gene expression, we focused on ASO1, a gapmer designed to target regRNAs transcribed from the Crick (minus) strand of the *OTC* enhancer. ASO1 consistently and specifically upregulated *OTC* mRNA and protein levels in PHHs (Figures 3C and 4A). Moreover, ASO1 exhibited no sequence complementarity to the alternatively spliced *OTC* mRNA isoform identified in PHHs and human liver tissue, facilitating a more direct interpretation of its mechanism of action.

**Figure 4.**
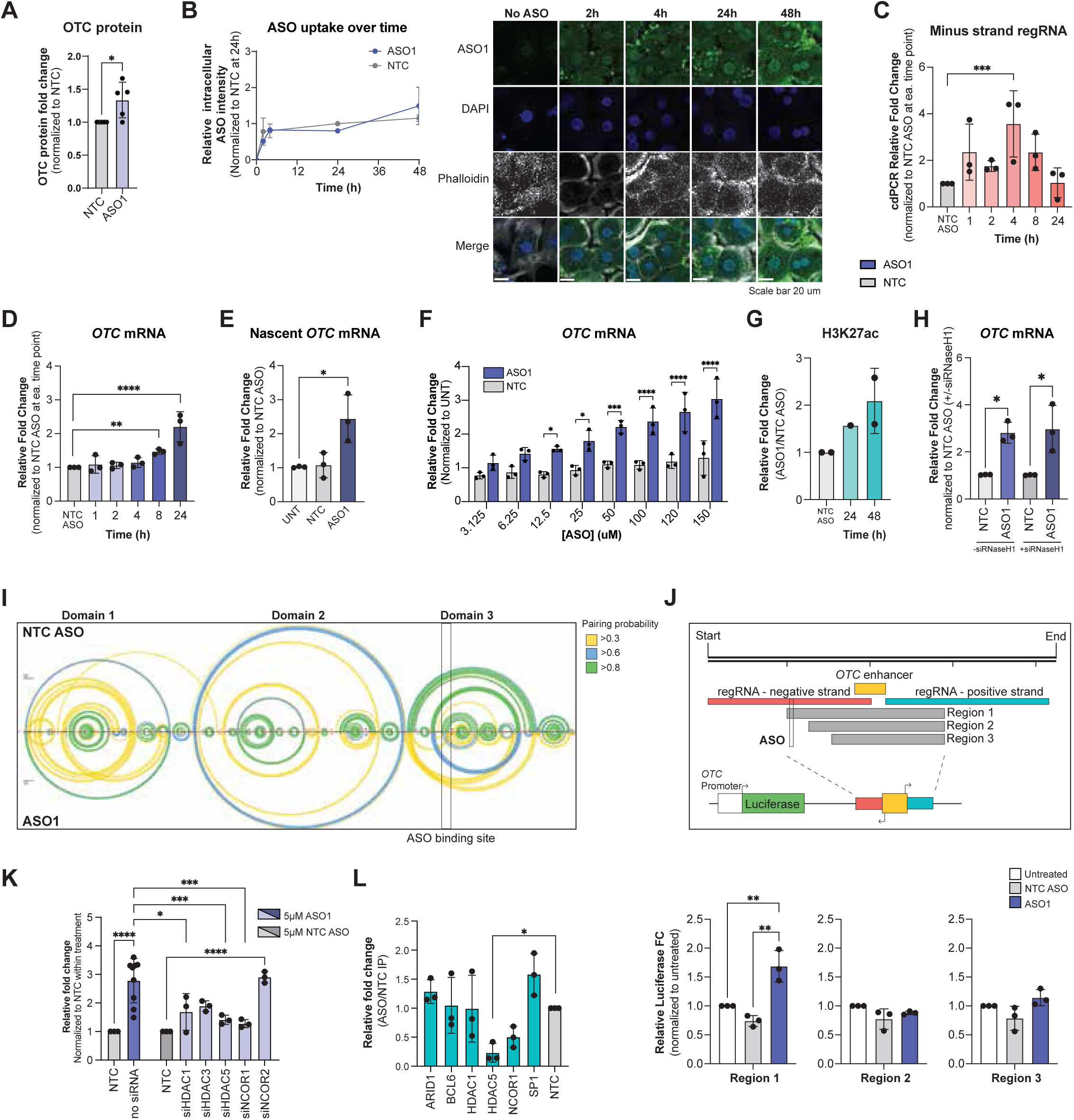
Mechanistic characterization of ASOs upregulating *OTC* expression in PHHs. (A) OTC protein measured by Jess^™^ capillary-based western blot system following treatment with ASO1. Protein data represent mean fold change relative to NTC ± s.d. (n = 5). Vinculin (VCL) was used as a loading control. Statistical significance was assessed using a paired *t*-test. (*, p<0.05) (B) Intracellular accumulation of ASO1 in PHHs over 48 h following gymnotic delivery (5 µM). Left, time course of ASO1 and NTC ASO accumulation in PHHs. Data are expressed as fold changes relative to NTC values ± s.d. (n = 2) measured at 24 h. Intracellular accumulation over time shows no significant difference between ASO1 and NTC treatments (p *=* 0.1536). Right, corresponding representative immunofluorescence images showing ASO1 accumulation over time. ASO1 was labeled with Alexa Fluor 488 dye via click chemistry, nuclei were stained with DAPI, and actin was detected with Phalloidin-Alexa Fluor 647 dye. (C) Expression of minus-strand *OTC* regRNA over 24 h following gymnotic delivery of ASO1 (5 µM), measured by cdRT-PCR. Data represent mean fold change relative to NTC ± s.d. (n = 3), normalized to telomerase RNA component (*TERC*). Statistical significance was determined by corrected unpaired *t*-test. (***, p<0.0007). (D) Time course of *OTC* mRNA upregulation following gymnotic delivery of ASO1 (5 µM) in PHHs over 24 h. ASO1 targets regRNAs transcribed from the minus strand of the *OTC* enhancer. mRNA data represent mean fold change relative to *PPIA* ± s.d. (n = 3). Statistical significance was assessed by corrected unpaired *t*-test. (**, p<0.005; ****, p<0.0001). (E) Nascent *OTC* transcript levels 48 h after treatment with ASO1 (5 µM). Untreated cells and those treated with a non-targeting control (NTC) ASO served as controls. Data are mean fold change ± s.d. (n = 3). Statistical analysis was performed using a paired *t*-test (*, p<0.025). (F) Dose-response of *OTC* mRNA expression following 48 h treatment with increasing concentrations of ASO1. Expression was normalized to NTC-treated cells at each concentration. Data are shown as mean fold change relative to *PPIA* ± s.d. (n = 3). Corrected unpaired *t*-test was used for statistical analysis. (*, p<0.035; ***, p<0.0007; ****, p<0.0001). (G) ChIP-qPCR analysis of H3K27Ac enrichment at the *OTC* enhancer 24 h and 48 h after ASO1 (5 µM) treatment. Data are presented as mean fold change relative to NTC ± s.d. (n = 3). (H) *OTC* expression 48 h after ASO1 (5 µM) treatment in PHHs pre-treated for 72 h with either control siRNA or siRNA targeting *RNASEH1* mRNA. Data represent mean fold change relative to NTC ± s.d. (n = 3), normalized to *PPIA*. Statistical significance was assessed by one-way ANOVA (*, p<0.018 and p<0.011 for ASO treatment without siRNA and with siRNA, respectively). (I) Linear arc diagram of nucleotide pairing probabilities in *OTC* regRNA, based on SHAPE-MaP data. Probabilities for ASO1-treated samples are shown below the x-axis, NTC-treated samples above. Boxed region indicates the ASO1 binding site. (J) Top, schematic of *OTC* enhancer regions used in a dual-luciferase reporter assay. The ∼4 kb enhancer locus was divided into three regions (1, 2, and 3) and cloned downstream of the *OTC* promoter-driven *Renilla* luciferase gene. Constructs were transfected into HepG2 cells. Bottom, relative Renilla luciferase activity (normalized to Firefly luciferase) following treatment with ASO1 (blue), NTC ASO (gray), or no ASO (white) for the three regions. Data represent fold change relative to no ASO ± s.d. (n = 3). Paired *t*-test was used for statistical comparison. (*, p<0.05). (K) ChIP-qPCR analysis of TF and co-repressor enrichment (ARID1, BCL6, HDAC1, HDAC5, NCOR1, and SP1) at the *OTC* enhancer 48 h after ASO1 (5 µM) treatment. Data are presented as mean fold change in enrichment relative to NTC ± s.d. (n = 3). Corrected paired *t*-test was used for statistical comparison. (*, p<0.0008). (L) *OTC* expression 48 h following ASO1 (5 µM) treatment in PHHs pre-treated with control siRNA or siRNAs targeting *HDAC1*, *HDAC3*, *HDAC5*, *NCOR1*, or *NCOR2* mRNAs for 72 h. Data represent mean fold change relative to NTC ± s.d. (n = 3), normalized to *PPIA*. Statistical significance was assessed with one-way ANOVA (*, p<0.02; ***, p<0.006; ****, p<0.0001).

To characterize the kinetics of ASO1 activity in PHHs, we tracked its intracellular uptake, effects on *OTC* regRNA and mRNA levels, and associated chromatin modifications at the *OTC* enhancer over time. ASO1 was delivered gymnotically at 5 µM and maintained in culture for up to 48 hours (h). An alkyne-modified ASO1, labeled post-fixation with an azide-conjugated fluorophore via click chemistry, was used to monitor intracellular accumulation. ASO1 was detectable within 2 h of treatment and continued to accumulate, reaching near-saturation by 24 h (Figures 4B and S5A).

Strand-specific crystal digital PCR (cdPCR) developed to detect changes in the steady-state levels of the minus-strand *OTC* regRNAs revealed extremely low baseline levels (∼0.22 molecules per cell) (Figure S5B). Following ASO1 treatment, regRNA levels rose transiently, peaking at 4 h, and declining by 24 h, despite continued intracellular accumulation of ASO1 (Figure 4B and 4C). In contrast, *OTC* mRNA levels increased progressively, with significant upregulation evident by 8 h, and exceeding a two-fold induction by 24 h (Figure 4D). These elevated levels were maintained at 48 h (Figure S5C), consistent with a sustained effect. Notably, *OTC* upregulation was accompanied by increased levels of nascent *OTC* transcripts (Figure 4E), indicating that ASO1 enhances gene expression at the level of transcription.

To assess the dose-responsiveness of this effect, ASO1 concentrations were titrated up to 150 µM, the highest possible under current hepatocyte culture conditions. *OTC* mRNA levels increased in a concentration-dependent manner, reaching up to a three-fold induction without evidence of saturation (Figure 4F), suggesting that both ASO1 concentration and exposure time contribute to the magnitude of gene upregulation.

To evaluate *OTC* enhancer activity directly, we examined changes in H3K27Ac, a canonical histone mark associated with active enhancers, at the enhancer. ChIP-qPCR revealed a time-dependent increase in H3K27Ac enrichment, with a two-fold elevation by 48 h following ASO1 treatment (Figure 4G).

Because ASO1 is a gapmer, we next tested whether its regulatory effects depend on RNase H1-mediated degradation of targeted regRNAs. siRNA-mediated knockdown of RNase H1 did not attenuate ASO1-induced *OTC* upregulation (Figures 4H and S5D), indicating that ASO1 acts independently of canonical RNase H1 activity.

These findings demonstrate a positive correlation between changes in regRNA expression and *OTC* mRNA induction in response to ASO1 treatment. They suggest that intracellular accumulation of ASO1 triggers a transient upregulation of enhancer-associated regRNAs, which precedes the more sustained upregulation of *OTC* mRNA and enhancer activation.

We hypothesized that ASO1 may enhance *OTC* transcription by inducing structural changes in the targeted enhancer-derived regRNAs which leads to increased local enhancer activity. To test this, we performed selective 2′-hydroxyl acylation analyzed by primer extension coupled with mutational profiling (SHAPE-MaP) *in vitro*, comparing structural changes elicited by ASO1 to those induced by a non-targeting control ASO. SHAPE-MaP analysis revealed three major structural domains within the regRNA, each comprising multiple subdomains formed through long-range intramolecular interactions (Figure 4I). ASO1 induced pronounced structural rearrangements both locally at its binding site and at distal regions (Figure 4I, bottom). Notably, these changes were predominantly localized to the third domain of the regRNA. In contrast, the control ASO did not induce local structural changes at the ASO1 binding site, nor did it affect SHAPE-MaP reactivity at distal regions (Figure 4I, top). Furthermore, sequence substitutions within the ASO1 binding site of the regRNA abolished these structural changes, indicating that an intact ASO1 binding site is essential for mediating the observed conformational rearrangements (Figure S5E).

To confirm that the ASO1 binding site is required for enhancer activation, we employed a dual-luciferase reporter assay. Regions of 1-2 kb encompassing the *OTC* enhancer, either containing (region 1) or lacking (regions 2 and 3) the ASO1 binding site, were cloned downstream of the *Renilla* luciferase gene under the control of the *OTC* promoter in a reporter plasmid and transfected into HepG2 cells (Figure 4J, top).

Reporter activity was quantified relative to a co-transfected *Firefly* luciferase control plasmid to normalize for transfection efficiency. A significant increase in luciferase activity was observed in response to ASO1, but only for constructs containing the ASO1 binding site (Figure 4J, bottom). Together, these results indicate that the ASO1 binding site is required for ASO1-mediated enhancer activation.

Given the established role of eRNAs and paRNAs in facilitating interactions with activating TFs^1,9^, and the likelihood that both activating and repressive TFs may compete for binding at regulatory elements, we hypothesized that ASO1 binding to the *OTC* regRNA alters this balance: reduces repressive TF occupancy and/or enhances recruitment of activating TFs and, thereby, promoting enhancer activation.

To test whether ASO1 affects the binding of repressive TFs and associated co-repressor complexes at the targeted enhancer, we curated a list of >200 proteins using available public data in HepG2 cells^54,55^ and refined it to six candidates with reported repressor activities. These candidates included two TFs (SP1^56,57^ and BCL6^58^), a component of the SWI/SNF chromatin remodeling complex (ARID1^59^), and three co-repressor complex components: histone deacetylase 1 (HDAC1^60^), which forms the catalytic core of multiple co-repressor complexes^61–64^, histone deacetylase 5 (HDAC5^65^), and nuclear receptor corepressor 1 (NCOR1^66^). All six candidates are expressed in PHHs and show significant enrichment at the *OTC* enhancer, based on publicly available ChIP-seq data from HepG2 cells (Figure S5F and Supplementary Table 7).

We next examined whether ASO1 binding to the *OTC* regRNA alters occupancy of the candidate repressors at the *OTC* enhancer. To this end, ChIP-qPCR was performed in PHHs using repressor-specific antibodies to assess binding of the six candidate repressors following ASO1 treatment. ASO1 treatment led to a significant reduction in HDAC5 binding at the *OTC* enhancer, with a modest decrease observed for NCOR1. No significant changes were detected in the binding of the remaining four repressors (Figure 4K). These findings suggest that reduced occupancy of HDAC5 and NCOR1 contributes to ASO1-mediated activation of the *OTC* enhancer.

To evaluate the functional contribution of these repressors to ASO1 activity, we conducted siRNA-mediated knockdown experiments to deplete each repressor in PHHs (Figure S5G). In addition to reducing levels of HDAC5 and NCOR1, we also decreased levels of histone deacetylase 3 (HDAC3^67^), an additional component of NCOR1 co-repressor complex, as well as NCOR2, a closely related co-repressor known to interact with HDAC3, and HDAC1. Partial depletion of each of these 5 repressors (*HDAC1*, *HDAC3*, *HDAC5*, *NCOR1, or NCOR2*) did not significantly increase *OTC* expression (Figure S5H). However, knockdown of *HDAC1*, *HDAC5*, or *NCOR1* attenuated ASO1-induced *OTC* mRNA upregulation, with *HDAC5* and *NCOR1* showing the most pronounced effects. In contrast, *NCOR2* knockdown had no effect on ASO1-induced *OTC* mRNA upregulation, while *HDAC3* knockdown resulted in nominal, but not statistically significant, reduction in ASO1-mediated effects (Figure 4L).

Together, these findings suggest that ASO1-mediated *OTC* mRNA upregulation requires normal levels of *HDAC1*, *HDAC5* and *NCOR1*, implicating these repressive factors as key mediators of ASO1-induced *OTC* mRNA upregulation. Our data support a model in which ASO1 binding induces structural changes in the targeted eRNA transcribed by the *OTC* enhancer, leading to eviction of transcriptional repressors from the enhancer and increased enhancer activity, ultimately driving upregulation of *OTC* gene expression.

## Discussion

In this study, we present a high-resolution catalog of regRNA species in PHHs, enabled by regRNA Capture-seq, a targeted long-read sequencing method optimized for detecting both polyadenylated and non-polyadenylated, low-abundance transcripts (Figure 1). The use of PHHs, a physiologically relevant yet technically challenging primary cell type, adds both biological and translational significance to our work. PHHs retain native metabolic and transcriptional programs often lost in immortalized liver-derived lines but have remained underutilized in eRNA studies due to technical limitations^68^. Our approach overcomes these barriers, providing the most comprehensive annotation of regRNAs in PHHs to date.

Integration of ATAC-seq and H3K27Ac ChIP-seq data identified over 134,000 active enhancers in PHHs, the majority of which were intragenic and transcriptionally active (Figures S1A and S1B). Using PRO-seq, we confirmed that eRNAs and paRNAs exhibited comparable transcriptional densities (Figure S1). These findings support the view that enhancers and promoters share many core transcriptional features, including RNA Pol II engagement and chromatin accessibility^69–71^. While bidirectional transcription predominated, a subset of enhancers produced eRNAs unidirectionally (Figure S2), potentially reflecting regulatory asymmetry, differential regRNA stability and turnover^72,73^, or limits of detection sensitivity for very low-abundance regRNAs.

By coupling targeted regRNA capture with long-read sequencing, we developed regRNA Capture-seq that offers high sensitivity, isoform-level resolution, and scalability, enabling detection of regRNAs from single enhancers or promoters as well as from thousands of regulatory elements in parallel. Unlike nascent RNA techniques such as PRO-seq, which capture transcriptional elongation with nucleotide precision but do not provide transcript-level resolution, regRNA Capture-seq reconstructs full-length isoforms, revealing splicing, polyadenylation, and transcript composition. Moreover, whereas PRO-seq identifies transient, unprocessed RNAs, regRNA Capture-seq enables detection of both nascent and processed regRNAs, providing a more complete picture of enhancer– and promoter-associated transcription.

Using this method, we identified over 19,000 distinct regRNAs arising from ∼2,000 enhancers in PHHs, 96.8% of which were previously unannotated (Figure 1E). Most regRNAs were unspliced and non-polyadenylated, consistent with properties of unstable nuclear transcripts. However, many enhancers also expressed longer, spliced, and polyadenylated isoforms. These observations point to unanticipated regRNA isoform diversity and suggest that individual enhancers may generate multiple functionally distinct regRNA isoforms.

On average, each enhancer generated approximately seven distinct regRNA transcripts, encompassing both polyadenylated and non-polyadenylated species (Figure S2E). Both RNA classes exhibited pronounced 3′ end heterogeneity (Figure S2I and S2J). For non-polyadenylated transcripts, this variability likely reflects a combination of ongoing nascent transcription and exonucleolytic processing by the RNA exosome complex^11,35^. In contrast, heterogeneity among polyadenylated regRNAs is consistent with the utilization of cryptic polyadenylation sites, which in canonical mRNAs are typically suppressed by U1 snRNP^74,75^.

Only 32.7% of regRNA species detected in PHHs were multi-exonic, and just 29.3% of these contained canonical splice motifs, indicating that fewer than 10% of regRNAs undergo conventional splicing (Table 1). Multi-exonic transcripts, whether harboring canonical motifs or not, were longer than unspliced species (Figure S2K), suggesting that they are not sequencing artefacts. Instead, these observations are consistent with recent evidence that lncRNAs generally exhibit weaker splice site motifs, lower splicing efficiency, and shorter introns than mRNAs, producing fewer fully spliced isoforms^76,77^. Together, our findings suggest that regRNAs are largely subject to minimal splicing constraints, which likely contributes to their rapid processing, limited half-life, and reduced alternative isoform diversity relative to mRNAs.

Although regRNA Capture-seq is inherently semi-quantitative due to the absence of unique molecular identifiers (UMIs) (Figure S2G), and extremely short or highly unstable RNAs may be identified less efficiently, the approach enables isoform-level resolution not possible for the short-read or nascent RNA methods. The strong concordance with PRO-seq (Figure S2D) supports its accuracy in detecting transcripts produced by active regulatory elements, while its full-length resolution allows dissection of their isoform-level complexity.

To further explore functional implications, we applied regRNA Capture-seq to a genetically validated enhancer regulating the *OTC* gene, the most frequently mutated gene in UCDs (Figure 2). Several lines of evidence confirmed transcriptional activity at this enhancer, including expression of multiple eRNA isoforms in PHHs and human liver tissue (Figures 2C, S3D, and S3E). Unexpectedly, we also identified spliced, polyadenylated transcripts initiating at the enhancer and extending into the *OTC* gene body (Figure S3E). These isoforms closely resembled canonical *OTC* mRNA but contained one or two upstream non-coding exons absent from the reference annotation (Figure S3F). These findings suggest that the *OTC* enhancer may also function as an alternative promoter, consistent with prior observations that enhancers can give rise to stable spliced RNAs and serve as alternative TSSs for coding genes^11,72,78^.

To evaluate the functional role of these regRNAs, we designed and screened ASOs targeting the *OTC* enhancer-derived transcripts in PHHs and evaluated their effect on *OTC* gene expression (Figure 3). Most ASOs showed no effect on *OTC* expression, while a small subset increased *OTC* mRNA levels in a concentration-dependent manner (Figures 3A and 3B). This is one of the first demonstrations that ASOs can be used to upregulate gene expression by selectively targeting eRNAs, representing a potentially new paradigm in RNA therapeutics^79,80^. Prior efforts have primarily focused on ASOs for transcript degradation or splice modulation^81^, whereas direct gene activation through regRNA targeting remained largely unexplored. The fact that only a small subset of regRNA-targeting ASOs upregulated *OTC* expression suggests that specific regions within the eRNA may play a key role in regulation of *OTC* gene expression. It is also possible that a precise combination of ASO sequence and chemistry might be required for effective engagement of the target eRNA and modulation of gene expression. Together, these observations highlight the need for future work to map functionally-relevant regions within regRNAs and underscore the importance of ASO design, consistent with the extensive body of prior work demonstrating effects of sequence and chemistry on ASO-mediated mRNA degradation and splice modulation^82–85^.

To begin to elucidate the mechanisms underlying *OTC* upregulation in response to *OTC* eRNA-targeting ASOs, we evaluated ASO-induced changes in *OTC* eRNA secondary structure and abundance as well as changes in TF occupancy and chromatin state at the targeted enhancer (Figure 4). ASO binding altered the secondary structure of the targeted eRNA *in vitro*, as shown by chemical probing (Figure 4I). These structural changes were associated with increased steady-state eRNA levels (Figure 4C) as well as elevated nascent mRNA levels in PHHs (Figure 4E). Concomitantly, we observed reduced binding of negative transcriptional regulators (Figure 4K) and increased H3K27 acetylation (Figure 4G) at the targeted enhancer indicative of elevated enhancer activity. Although SHAPE-based chemical probing was limited to cell-free analysis due to the extremely low expression of *OTC* regRNAs in PHHs (fewer than 0.3 molecules per cell), future advances in chemical probing sensitivity are expected to enable in-cell analysis.

Together, these findings support a model, in which ASO binding induces structural changes in the targeted eRNA, promoting eviction of transcriptional repressors from the enhancer, and potentiating enhancer function to increase gene transcription. Importantly, we cannot exclude the possibility that ASOs also act by stabilizing the eRNA or enhancing TF recruitment to the enhancer. It remains unclear whether increased eRNA levels precede enhancer activation or result from it. These possible alternative mechanisms underscore the complexity of ASO-mediated eRNA modulation and highlight the need for further studies incorporating in-cell structural profiling and time-resolved analyses.

While ASOs have been reported to induce RNA degradation or exert off-target effects in some settings^51,86^, our data suggest these mechanisms are unlikely to explain the observed effects on *OTC* expression. Since ASO-induced upregulation did not require RNase H1 activity, even for a gapmer ASO (Figure 4H), the results are more consistent with ASO-induced stabilization or structural remodeling of the targeted eRNA. This suggests that, despite its gapmer design, the ASO predominantly acts through steric hindrance rather than by promoting target eRNA cleavage. Mechanistically, these findings align with previous studies showing that non-coding RNAs can influence chromatin accessibility and TF dynamics via RNA secondary structure^13,71,87^, and that ASO binding can modulate RNA-protein interactions to alter gene expression^88,89^.

In summary, regRNA Capture-seq enables high-resolution mapping of regRNAs in primary human cells and reveals unanticipated isoform complexity and regulatory capacity. Our identification of the *OTC* enhancer with dual roles, as both a source of eRNAs and an alternative promoter, illustrates the architectural and functional complexity of enhancer transcription. The demonstration that ASOs can upregulate gene expression *in vitro* by targeting eRNAs potentially introduces a novel therapeutic strategy for modulating gene activity via non-coding RNA elements. These findings lay a foundation for the mechanistic dissection and therapeutic targeting of regRNAs in human physiology and disease.

## Figure Legends

**Supplementary Figure 1.** Characterization of active enhancers in primary human hepatocytes. (A) Venn diagram showing overlap between ATAC-seq summits and H3K27Ac peaks (±500 nt), used to define 134,520 active enhancers in PHHs. (B) Genomic classification of hepatocyte enhancers relative to position of protein-coding genes (GENCODE v43). (C) Heatmaps of PRO-seq (left), H3K27Ac ChIP-seq (center), and ATAC-seq (right) signal at active enhancers, centered on ATAC-seq summits (± 1,500 nt). Color scale represents normalized read coverage (RPM) per 10 nt bin. Signal above the dashed line corresponds to the 96,169 enhancers exhibiting detectable transcriptional activity, defined as >0.02 PRO-seq reads per million (RPM). (D) Histogram of PRO-seq signal intensities (RPM) for the top 10,000 paRNA– and eRNA-producing regions. PRO-seq signal was measured within a 1 kb window centered on ATAC-seq peak summits. Reads mapped to the plus and minus strands were quantified separately. For enhancers (not overlapping promoters), PRO-seq signal was measured within ±500 nt of ATAC-seq peak summits. For intragenic enhancers, only antisense signal relative to the direction of transcription of the corresponding host gene was quantified. PRO-seq signals for paRNAs were similarly quantified within ±500 nt of ATAC-seq peak summits located within ±1 kb of annotated TSSs using only antisense signal relative to gene transcription. Mean and median values are indicated. (E) Histogram showing PRO-seq read distribution for 13,979 paRNA– and 68,052 eRNA-producing loci. PRO-seq signals were calculated as in (D). Intragenic enhancers with only sense PRO-seq signal relative to the host gene transcription were excluded. Mean and median values are indicated.

**Supplementary Figure 2.** Characterization of regRNAs in the regRNA catalog in PHHs. (A) Boxplot of PRO-seq signal (RPM) across four stratified enhancer groups used in the regRNA Capture-seq in PHHs. (B) Genomic classification of ∼2,000 probed enhancers relative to position of protein-coding genes (GENCODE v43). (C) Heatmaps of polyadenylated Capture-seq, non-polyadenylated Capture-seq, and PRO-seq reads centered on enhancer ATAC-seq summits (±5 kb) for the four stratified enhancer groups. Color bar scale represents normalized read counts per 10 nt bin. (D) Mean signal profiles of Capture-seq and PRO-seq reads relative to enhancer center (±5 kb). (E) Histogram showing the number of regRNAs per enhancer. Median is indicated. (F) Number of captured polyadenylated (polyA) and non-polyadenylated (non-polyA) eRNA and paRNA species across the four stratified enhancer groups. (G) Boxplot of Capture-seq signal (RPM) supporting individual regRNA annotations across four stratified enhancer groups in PHHs. (H) Histogram showing distribution of lengths (nt) of captured polyadenylated (polyA) and non-polyadenylated (non-polyA) regRNAs. Median lengths are indicated. (I) Distance from 5′ and 3′ ends of polyadenylated (polyA) mono– and multi-exonic regRNAs to putative isoform boundaries. Paired two-sided Student’s *t*-test used. (J) Same as in (I), for non-polyadenylated (non-polyA) mono– and multi-exonic regRNAs. (K) Histogram of length distributions for polyadenylated (polyA) and non-polyadenylated (non-polyA) mono– and multi-exonic regRNAs with and without canonical splicing motifs. Median lengths are indicated. (L) Boxplot of regRNA read counts (RPM) for bidirectional enhancers, unidirectional enhancers, and promoters. Sample sizes are indicated above.

**Supplementary Figure 3.** The *OTC* enhancer is a transcriptionally active regulatory element across cell types. (A) Motif analysis centered on *OTC* enhancer defined by ATAC-seq summit (±250 nt). Colors denotes specific transcription factor motifs defined by Transfac database. (B) ChIP-seq signals for H3K4me3, H3K27Ac, and H3K4me1 at the *OTC* locus in HepG2 cells and PHHs. Signal intensity is shown in RPM. (C) Top, longest putative peptide lengths (x-axis) corresponding to the longest regRNA sequences translated in all three reading frames from the minus (left) and plus (right) strands of the *OTC* enhancer, shown relative to distributions of lengths for annotated coding and non-coding transcripts. Bottom, corresponding Fickett Testcode scores (x-axis) for the same regRNAs. (D) Positions of probes used for regRNA Capture-seq at the *OTC* enhancer shown relative to regRNA Capture-seq, ATAC-seq, and PRO-seq profiles in PHHs and human liver tissue. regRNA reads mapping to the plus strand are shown in blue, minus strand in red. *OTC* mRNA reads are excluded. PRO-seq reads mapping to the plus strand are shown above the x-axis, minus strand below. Signal intensity is shown in RPM. (E) regRNA Capture-seq, ATAC-seq, and PRO-seq profiles in PHHs and human liver tissue as shown in (C) but include Capture-seq reads for alternatively spliced *OTC* mRNA isoforms. (F) Abundance of extended alternatively spliced *OTC* mRNA isoforms measured by qRT-PCR shown relative to the abundance of the plus-strand *OTC* regRNA (left) and total *OTC* mRNA (right) in PHHs and human liver. Individual data points are from three different plateable PHHs donors and four different human donor liver tissue biopsies. Reverse transcription was performed using random primers (RP) or oligo(dT) primers (dT). (G) Volcano plot of Log_10_P-values (Log_10_P) (y-axis) relative to Log_2_ fold-changes in gene expression (Log_2_FC) (x-axis) measured by RNA-seq in PHHs treated with JAKi. Significantly downregulated and upregulated genes are colored in green and pink respectively, with *OTC* highlighted in red.

**Supplementary Figure 4.** Identification and optimization of ASOs upregulating *OTC* gene expression in PHHs. (A) *OTC* mRNA levels measured by qRT-PCR in PHHs treated with 5 µM ASOs targeting either minus-strand regRNAs (left) or plus-strand regRNAs (right). Data correspond to Figure 3A, shown here separately for gapmer and steric-blocking ASO designs. Untreated cells and cells treated with three non-targeting control (NTC) ASOs served as controls. Data represent mean fold change relative to *B2M* ± s.d. (n = 3). Statistical significance was assessed by one-way ANOVA. (B) Dose-response curves showing changes in mRNA levels of retinitis pigmentosa GTPase regulator (*RPGR*), *OTC* neighboring gene, measured by qRT-PCR following treatment of PHHs with increasing concentrations of the five *OTC* regRNA-targeting ASOs shown in Figure 3C. Untreated and NTC ASO-treated cells served as controls. Data are shown as mean fold change relative to *B2M* ± s.d. (n = 3). ASO designs are illustrated above the graphs. Black circles represent 2′-MOE-modified nucleotides, white circles denote DNA nucleotides. (C) Left, ASO65-ASO75 positions are shown relative to the positions of the targeted plus-strand *OTC* regRNAs. Right, *OTC* mRNA levels measured by qRT-PCR in PHHs treated with 5 µM ASOs targeting the plus-strand *OTC* regRNA. Data are shown as mean fold change relative to *B2M* ± s.d. (n = 3). (D) Dose-response curves showing *OTC* mRNA levels following treatment of PHHs derived from two different donors with increasing concentrations of ASO80 and ASO83 shown in Figure 3D, targeting either minus– (red) or plus-strand (blue) *OTC* regRNAs. Untreated and NTC ASO-treated cells served as controls. Data are shown as mean fold change relative to *B2M* ± s.d. (n = 3).

**Supplementary Figure 5.** Mechanistic characterization of ASOs upregulating *OTC* expression in PHHs. (A) Intracellular accumulation of NTC ASO in PHHs over 48 h following gymnotic delivery (5 µM). NTC was labeled with Alexa Fluor 488 dye via click chemistry, nuclei were stained with DAPI, and actin was detected with Phalloidin-Alexa Fluor 647 dye. (B) Left, standard curve used to estimate the number of *OTC* regRNA copies per µl (y-axis) based on the number of positive droplets/µl (x-axis) measured by cdPCR using serial dilutions of *in vitro*-transcribed *OTC* regRNA and an *OTC* regRNA-specific probe. Right, cdPCR analysis of serial dilutions of total RNA isolated from PHHs, used to quantify the number of positive droplets/µl (y-axis) and estimate regRNA copies per cell (see Methods). (C) *OTC* mRNA levels measured by qRT-PCR in PHHs treated with 5 µM ASO1 for 48 h. Data represent mean fold change relative to NTC ± s.d. (n = 3), normalized to *PPIA*. Statistical significance was assessed using a paired *t*-test (*, p<0.05). (D) *RNASEH1* expression in PHHs following 72 h knockdown with gene-specific siRNAs. Expression was normalized to untreated cells at the same time points. Data are shown as mean fold change relative to *GAPDH* ± s.d. (n = 3). Statistical analysis was performed using one-way ANOVA. (***, p<0.0001) (E) Top, heatmap showing changes in SHAPE-MaP reactivity in *OTC* regRNA following treatment with ASO1 compared to a non-targeting control (NTC) ASO. Bottom, heatmap showing ASO1-induced reactivity changes in *OTC* regRNA bearing a mutated ASO1-binding site. Arrows indicate positions complementary to ASO1. (F) Signal tracks for ATAC-seq shown alongside ChIP-seq data for H3K27Ac, SP1, ARID1A, BCL6, NCOR1, and HDAC1 at the *OTC* locus in HepG2 cells. Signal intensity is shown in RPM. (G) Expression of *HDAC1*, *HDAC3*, *HDAC5*, *NCOR1*, and *NCOR2* in PHHs following 72 h knockdown with gene-specific siRNAs. Expression levels are normalized to untreated cells. Data are presented as mean fold change relative to *GAPDH* ± s.d. (n = 3). Statistical significance was assessed using one-way ANOVA (****, p<0.0001). (H) Expression of *OTC* in PHHs following 72 h knockdown with gene-specific siRNAs targeting the same genes as in (G). Expression levels are normalized to untreated cells. Data are presented as mean fold change relative to *GAPDH* ± s.d. (n = 3). Statistical significance was assessed using one-way ANOVA (ns).

## Methods

### Resource availability

Requests for further information and resources should be directed to and will be fulfilled by the lead contact, Alla A. Sigova.

## Materials availability

All requests for materials will be promptly reviewed by CAMP4 Therapeutics Corp. to verify whether the request is subject to any intellectual property or confidentiality obligations. Following review, materials can be shared via a material transfer agreement by contacting the lead contact.

## Data availability

- Information for publicly available datasets used can be found in Supplementary Table 7

**Primary human hepatocyte culture.** Cryopreserved primary human hepatocytes (PHHs) collected under informed consent (#HUPCI, Lonza) were thawed in a 37 °C water bath for 2 minutes (min), then transferred to 50 mL of Human Cyropreserved Hepatocyte Thawing Medium (#MCHT50, Lonza). Cells were centrifuged at 100 × g for 8 min at room temperature. The supernatant was aspirated, and the cell pellet was gently resuspended in Hepatocyte Plating Medium with supplement (#MP250, Lonza) and plated according to the manufacturer’s recommended seeding densities. Cells were maintained in a humidified incubator at 37 °C with 5% CO₂. For regRNA measurements, hepatocytes were seeded at a density of 5.5 × 10 cells/well in 12-well collagen-coated plates (#354500, Corning Inc.). After 4 hours, plating medium was aspirated and replaced with Cellartis^®^ Power^™^ Primary HEP Medium (#Y20020, Takara Bio USA, Inc.). Media was refreshed every 2 days for up to 6 days in culture. Cells were treated with 5 µM ASO for gymnotic delivery at designated time points.

**siRNA-based knockdowns in primary human hepatocytes**. PHHs (were seeded at 1.5 × 10 cells per well in 48-well collagen-coated plates (#354505, Corning Inc.). Three days post-plating, cells were transfected with 10 nM siRNA (Supplementary Table 6; Horizon Discovery) using Lipofectamine^™^ RNAiMAX Transfection Reagent (#13778150, Thermo Fisher Scientific Inc.), following the manufacturer’s instructions. 24 hours later, medium was replaced with fresh medium containing 5 µM ASO. Cells were harvested 48 hours after ASO treatment (6 days total in culture). Total RNA was extracted using the MagMAX^™^ mirVana^™^ Total RNA Isolation Kit (#A27828, Thermo Fisher Scientific Inc.), and 10 µL of RNA was used for cDNA synthesis using the SuperScript^™^ IV Reverse Transcriptase (#18090050, Thermo Fisher Scientific Inc.). *OTC* expression levels were quantified using commercially available quantitative real-time PCR (qRT-PCR) assays (Supplementary Table 2).

**Small molecule treatment of primary human hepatocytes**. PHHs were plated at 1.5 × 10 cells/well in 48-well collagen-coated plates. On day 5 post-plating, cells were treated with either 3 µM or 10 µM upadacitinib (#HY-19569, MedChemExpress) (for PRO-seq or *OTC* expression assays, respectively). RNA extraction, cDNA synthesis, and *OTC* quantification were performed as described above.

**dCas9-VPH and dCas9-KRAB lentivirus production**. Lentiviral constructs encoding either dCas9-VPH (activator) or dCas9-KRAB (repressor) were packaged by transfecting 9 × 10 Lenti-X^™^ 293T cells (#632180, Takara Bio USA Inc.) with 800 ng of CMV:dCas9-VPH-2A-Blast (#SVVPHC9B-PS, Cellecta, Inc.) or CMV:dCas9-KRAB-2A-Blast (#SVKRABC9B-PS, Cellecta, Inc.), 800 ng pVSVG, and 4 µg psPAX2 packaging plasmid, using FuGENE^®^ 6 Transfection Reagent (#E2692, Promega Corp.). Media was replaced 24 hours post-transfection. 72 hours post-transfection, viral supernatant was collected, filtered through a 0.45 µm polyethersulfone filter, and stored at –80 °C.

**Generation of HepG2-dCas9 cell lines.** HepG2 cells (#HB-8065, ATCC) were seeded at 2.5 × 10 cells per well in a 24-well plate and transduced at a high MOI with dCas9-VPH or dCas9-KRAB lentivirus and LentiTrans^™^ Transduction Reagent (#LTDR1, Cellecta, Inc.). Transduced cells were selected with 10 µg/mL blasticidin and expanded to generate stable HepG2-dCas9-VPH and HepG2-dCas9-KRAB cell lines.

**Single-guide RNA (sgRNA) design, cloning, packaging, and transduction.** sgRNAs targeting the *OTC* transcription start site (TSS) and putative enhancer regions were designed and cloned into a modified CROP-seq vector^90^ via *BsmBI* restriction sites (Supplementary Table 1). Lentiviral constructs encoding the sgRNAs were packaged in Lenti-X 293T cells as described above. Stable HepG2-dCas9-VPH and HepG2-dCas9-KRAB cell lines were transduced at a high MOI with sgRNA-expressing lentivirus and LentiTrans^™^ Transduction Reagent. Transduced cells were selected with 1 µg/mL puromycin to enrich for sgRNA-positive cells. Cells were harvested 5 days post-sgRNA transduction. Total RNA was extracted as described above. cDNA was synthesized as described above, and *OTC* expression was measured using qRT-PCR assays (Supplementary Table 2). Data were visualized using Prism software version 10.6.0 (GraphPad Software) with expression values plotted as fold change (FC) mean ± standard deviation (SD) relative to non-targeting control.

**ASO design and synthesis.** ASOs were designed as 20-mers tiled across the full length of *OTC* regRNAs, excluding highly repetitive regions and sequences with fewer than three nucleotide mismatches to other transcribed genomic loci. ASOs were synthesized either as gapmers with 2′-O-methoxyethyl (2′-MOE) modifications on the five terminal nucleotides at each end and a central DNA core or as fully 2’-MOE-modified oligonucleotides. All ASOs incorporated full phosphorothioate backbones to enhance nuclease resistance and RNA-binding affinity. Oligonucleotides were synthesized using solid-phase synthesis and purified by high-performance liquid chromatography (HPLC).

**ASO screening in primary human hepatocytes.** PHHs were seeded at 5.5 × 10 cells per well in 96-well collagen-coated plates (#354650, Corning Inc.). At 4 hours post-plating, cells were treated with 5 µM ASO via gymnosis in Cellartis Power Primary Hepatocyte Medium. After 42 hours of ASO treatment, cells were harvested and total RNA was extracted using the RNeasy^®^ 96 Kit (#74182, Qiagen N.V.) and eluted in 50 µL of RNase-free water. For cDNA synthesis, 10 µL of total RNA was reverse transcribed using the High-Capacity cDNA Reverse Transcription Kit (#4368813, Thermo Fisher Scientific Inc.). Changes in *OTC* gene expression were quantified using commercially available qRT-PCR assays (Supplementary Table 2).

**Quantitative reverse transcription followed by PCR (qRT-PCR) analyses.** Fold changes in gene expression between experimental and control treatments were calculated using a ΔΔCT approach^91^ as specified in Figure Legends and Supplementary Tables 2 and 5. Statistical analyses were conducted using Prism software version 10.6.0.

**regRNA Capture-seq in primary human hepatocytes.** PHHs were cultured for 6 days as described above. Cells were briefly washed 2X with 1 mL of phosphate-buffered saline (PBS) and total RNA extracted using TRIzol^™^ Reagent (#15596026, Thermo Fisher Scientific Inc.) following manufacturer’s recommendations. RNA was DNase I-treated and concentrated using the RNA Clean & Concentrator-25^™^ kit (#R1017, Zymo Research Corp.). Total RNA was separated into polyadenylated and non-polyadenylated RNA using previously published methodology^92^, with the exception that poly(A)-tailing was performed using *E. coli* poly(A) polymerase (#M0276S, New England Biolabs, Inc.). Double-stranded cDNA was PCR-amplified using Platinum^™^ SuperFi II PCR Master Mix (#12368010, Thermo Fisher Scientific Inc.). For regRNA catalog generation, 2,000 enhancers were stratified into four groups based on PRO-seq signal (very high/high/medium/low) and four corresponding probe sets, each targeting a panel of 500 different enhancers, were designed and synthesized (myBaits^®^ Custom Hybridization Capture Kit; Daicel Arbor Biosciences). Equal amounts of cDNA (>300 ng) were generated from both polyadenylated and non-polyadenylated RNA and used as input material for target enrichment using each of the four probe sets. regRNA capture was performed according to the probe manufacturer’s recommendations. For *OTC* regRNA Capture-seq, sequences of *OTC* enhancer and additional liver-specific control gene regions were used for probe design and synthesis. Long-read libraries were prepared using captured double-stranded cDNA with the SMRTbell^®^ Express Template Prep Kit 2.0 (#100-938-900, Pacific Biosciences of California, Inc.) according to manufacturer’s instructions, and sequenced on a Sequel^®^ II instrument (Pacific Biosciences of California, Inc.).

**Assay for transposase-accessible chromatin using sequencing (ATAC-seq).** ATAC-seq libraries were generated as previously described^93^, with minor modifications. Briefly, 5 x10^4^ PHHs were harvested after 6 days in culture, washed twice with cold 1× PBS, and lysed in cold lysis buffer (10 mM Tris-HCl, pH 7.4; 10 mM NaCl; 3 mM MgCl₂; and 0.1% octylphenoxypolyethoxyethanol for 10 min on ice to isolate nuclei.

Transposition reactions were performed at 37°C for 30 min using the Tn5 transposase from the Nextera^®^ DNA Library Prep Kit (#FC-121-1030, Illumina, Inc.). Transposed DNA was purified using the MinElute^™^ PCR Purification Kit (#28004, Qiagen N.V.) and amplified by PCR with indexed primers. Libraries were size-selected to enrich for fragments <1 kb using AMPure XP beads and quantified by qPCR. ATAC-seq libraries were sequenced on a HiSeq X^®^ system (Illumina, Inc.) to generate 150-bp paired-end reads.

**Chromatin immunoprecipitation followed by sequencing (ChIP-seq).** PHHs were plated on 100 mm collagen-coated dishes (#354450, Corning Inc.) at a density of 8 × 10^6^ cells per plate and cultured for six days, as described above. Cells were crosslinked in 1% formaldehyde (final concentration) for 15 min at room temperature, quenched with 2.5 M glycine for 5 min, and washed twice with ice-cold 1× PBS (#10010023, Gibco). Cells were scraped into 15 mL conical tubes, pelleted at 1,500 rpm for 5 min at 4°C, washed once more in 1× PBS, and flash-frozen at –80°C. Frozen crosslinked pellets were thawed on ice and lysed sequentially in three buffers (Buffers 1 and 2 and sonication buffer, described below), each supplemented with cOmplete^™^ Protease Inhibitor Cocktail (#04693116001, Roche). Composition of Buffer 1 was 50 mM HEPES, 140 mM NaCl, 1 mM EDTA, 10% glycerol, 0.5% octylphenoxypolyethoxyethanol, 0.25% Triton^®^ X-100; Buffer 2: 10 mM Tris-HCl (pH 8.0), 200 mM NaCl, 1 mM EDTA, 1 mM EGTA; and sonication buffer: 50 mM HEPES, 140 mM NaCl, 1 mM EDTA, 1% Triton X-100, 0.2% sodium deoxycholate, and 0.1% sodium dodecyl sulfate (SDS). Chromatin was sheared using a E220*evolution* Focused-ultrasonicator (Covaris, LLC). Lysates were centrifuged, and supernatants containing sheared chromatin were incubated overnight at 4°C with rotation with Dynabeads^®^ Protein G beads (#10004D, Thermo Fisher Scientific Inc.) pre-bound to antibodies (see Supplementary Table 4). Beads were washed extensively at 4°C with rotation, and immunoprecipitated material was eluted in buffer containing 50 mM Tris-HCl (pH 8.0), 10 mM EDTA, and 1% SDS at 65°C for 1 hour. Crosslinks were reversed by overnight incubation at 65°C. Immunoprecipitated and input DNA samples were treated with RNase A and Proteinase K, purified using MaXtract^™^ High Density Tubes (#129056, Qiagen N.V.), and ethanol precipitated overnight at –80°C. Sequencing libraries were prepared with the NEBNext^®^ Ultra^™^ II DNA Library Prep Kit for Illumina^®^ (#E7645, New England Biolabs, Inc.), following the manufacturer’s instructions, and sequenced on a HiSeq X system (Illumina, Inc.) to generate 150-bp paired-end reads.

**Precision run-on sequencing (PRO-seq).** PRO-seq libraries were generated using previously published protocols^94^. Briefly, PHHs were cultured as described above and harvested after 6 days, and *Drosophila* S2 cells (10% by cell number) were spiked-in, and permeabilized. Nuclear run-on was performed immediately at 37°C for 3 min in the presence of 0.5% sarkosyl, biotin-11-CTP (#NEL542001EA, Revvity Health Sciences, Inc.), and biotin-11-UTP (#NEL543001EA, Revvity Health Sciences, Inc.). RNA was fragmented and ligated with RNA adapters compatible with paired-end sequencing and UMI-specific data processing. Run-on transcripts were enriched over pre-existing RNA through three rounds of biotin-based pulldown at distinct stages of library preparation. Libraries were sequenced on a HiSeq X system (Illumina, Inc.) to generate 150-bp paired-end reads.

**Selective 2**′**-hydroxyl acylation analyzed by primer extension and mutational profiling (SHAPE-MaP).** Secondary structure of the *OTC* regRNA was assessed using SHAPE-MaP. The regRNA sequence (∼1,200 nt) was synthesized as a double-stranded DNA fragment (Supplementary Table 5) and directionally cloned downstream of the SP6 promoter of a high-copy vector designed for *in vitro* transcription. The plasmid was linearized with a restriction enzyme and used as template for *in vitro* transcription with the HiScribe^®^ SP6 RNA Synthesis Kit (#E2070S, New England Biolabs, Inc.), following the manufacturer’s protocol. Transcribed RNA was purified, treated with DNase I, and concentrated using the RNA Clean & Concentrator-25^™^ kit. For SHAPE labeling, 500 ng of purified RNA was denatured and refolded in the presence or absence of ASOs at a 5:1 ASO:RNA molar ratio. SHAPE reactions were performed at 37°C for 5 min in the presence of 20 mM 1-methyl-7-nitroisatoic anhydride (1M7) (#hy-d0913-50mg, MedChemExpress LLC). Each condition included a treated sample (+1M7), an untreated control (-1M7; DMSO only), and a denatured control. For the denatured control, RNA was incubated at 95°C for 1 minute in denaturation buffer (50 mM HEPES pH 8.0, 4 mM EDTA, 50% formamide) prior to SHAPE labeling. Reverse transcription was performed using SuperScript^™^ II reverse transcriptase (#18064014, Thermo Fisher Scientific Inc.) and random primers (#S1254S, New England Biolabs, Inc.) in a custom buffer containing 50 mM Tris-HCl (pH 8.0), 75 mM KCl, 6 mM MnCl₂, 10 mM dithiothreitol, and 0.5 mM dNTPs. cDNA was subjected to Tn5-based library preparation and amplified with seven cycles of PCR. Libraries were sequenced on an NovaSeq^®^ 6000 sequencing system (Illumina, Inc.) to a depth >5 million paired-end reads per sample.

**Chromatin immunoprecipitation followed by quantitative PCR (ChIP-qPCR).** ChIP-qPCR was performed using chromatin prepared as described above for ChIP-seq. Briefly, chromatin from PHHs was crosslinked with 1% formaldehyde, lysed, and sheared by sonication using a E220*evolution* Focused-ultrasonicator. Chromatin was incubated overnight at 4°C with rotation with Dynabeads Protein G beads pre-bound to the indicated antibodies (see Supplementary Table 4). Following immunoprecipitation, beads were washed extensively, and bound chromatin was eluted in buffer containing 50 mM Tris-HCl (pH 8.0), 10 mM EDTA, and 1% SDS at 65°C for 1 hour. Crosslinks were reversed by overnight incubation at 65°C. Immunoprecipitated and input DNA samples were treated with RNase A and proteinase K (#EO0491, Thermo Fisher Scientific Inc.), purified using MaXtract High Density Tubes, and ethanol precipitated overnight at –80°C. DNA was quantified and qPCR was performed using PowerUp^™^ SYBR^™^ Green PCR Master Mix for qPCR (#A25742, Thermo Fisher Scientific Inc.) on a QuantStudio^™^ 6 Pro Real-Time PCR System (Thermo Fisher Scientific Inc.). Primer sequences for target and control loci are listed in Supplementary Table 5. Enrichment was calculated as percent input or normalized to a reference region using the ΔΔCt method. Data visualization was performed using Prism software version 10.6.0 with expression values plotted as mean FC ± SD.

**RNA-seq.** PHHs were plated and cultured for 6 days as described above in 48-well collagen plates at a density of 1.5 × 10 cells/well. Cells were lysed and collected for RNA extraction as described above. 500 ng of total RNA was used to make RNA-seq libraries with the NEBNext Ultra Directional RNA Library Prep Kit for Illumina (#E7420, New England Biolabs, Inc.). Each library was made with a unique index of the NEBNext^®^ Multiplex Oligos for Illumina (#E6609, New England Biolabs, Inc.) and RNA-seq libraries were pooled and sequenced on a HiSeq X system (Illumina, Inc.) to generate 150-bp paired-end reads.

**Crystal digital PCR (cdPCR).** PHHs were cultured and treated for 6 days as described above. Cells (5.5 x 10) were lysed with TRIzol Reagent and RNA was isolated using manufacturer’s recommendations. Total RNA was DNase I treated and concentrated using the RNA Clean & Concentrator-25 kit (#R1017, Zymo). A total of 5 µg of RNA was used for cDNA synthesis per sample. 3 µL cDNA was combined with 3 µL mastermix containing naica^®^ PCR mix (#R10105, Stilla Technologies Inc.) and regRNA detection assays. A total of 5 µL of cDNA with mastermix was added to a single well of the Ruby Chip (#C16011, Stilla) and cDNA was amplified with 45X cycles to ensure sufficient number of positive regRNA droplets were detected using the Nio^®^ System (Bio-Rad Laboratories, Inc.). Custom expression detection assays were used to quantify regRNAs (Supplementary Table 5).

***OTC* enhancer reporter assay.** Various *OTC* enhancer constructs were PCR-amplified from human genomic DNA and cloned into the pGL4.10[*luc2*] vector (#E6651, Promega Corp.) downstream of the *OTC* promoter driving firefly luciferase expression. Constructs were verified by Sanger sequencing. For reporter assays, HepG2 cells were transfected with 500 ng of each enhancer-luciferase plasmid along with 50 ng of *Renilla* luciferase control reporter vector (pRL-TK, #E2241, Promega Corp.) as a transfection control, using Lipofectamine^™^ 3000 Transfection Reagent (#L3000015, Thermo Fisher Scientific Inc.) in 24-well plates. Dual-luciferase assays were performed 24-48 hours post-transfection as described above. Firefly luciferase activity was normalized to Renilla luciferase and reported relative to vector-only controls.

**OTC protein quantification.** For protein quantification in cultured cells, PHHs were cultured and treated as described above. Cells were washed once with warm PBS and lysed in RIPA buffer supplemented with Halt^™^ Protease and Phosphatase Inhibitor Cocktail (#PI78447, Fisher Scientific) for 15 min at 4°C. Lysates were briefly sonicated, centrifuged at 10,000 × *g* for 10 min at 4°C, and supernatants collected. Total protein concentration was measured using the Bicinchoninic Acid (BCA) Protein Assay Kit (#ab102536, Abcam Limited), to ensure equal loading. OTC protein levels were quantified using the Jess^™^ Automated Western Blot System (#004-650, Bio-Techne Corp.) using the 12-230 kDa Separation Module (#SM-W004, Bio-Techne Corp.) following the manufacturer’s protocol. Primary antibodies are listed in Supplementary Table 4. Protein expression was normalized to loading controls as indicated. Values were plotted in Prism software version 10.6.0 and shown as mean FC ± SD, relative to NTC ASO.

**Sample preparation for analysis of ASO localization.** PHHs were seeded into collagen I-coated, glass-bottom 96-well plates (#152036, Thermo Fisher Scientific Inc.) and cultured and treated as described above using alkyne-modified ASOs. After 48 hours, cells were fixed with 4% paraformaldehyde in PBS for 20 min at room temperature, washed three times with 1× PBS, and permeabilized in 1× PBS containing 0.1% Triton X-100 for 10 min. Following three additional 1× PBS washes, cells were incubated with TrueBlack^®^ Lipofuscin Autofluorescence Quencher (#23007, Biotium, Inc.) for 30 seconds to reduce background fluorescence and washed again three times with PBS. Cells were blocked overnight at 4°C in PBS containing 1% goat serum, 2% bovine serum albumin (BSA), and 0.02% polysorbate 20. The next day, cells were washed three times with PBS containing 0.02% polysorbate 20 and then three times with PBS containing 3% BSA. ASO labeling was performed using the Click-iT^™^ Plus Alexa Fluor^™^ 488 Picolyl Azide Toolkit (#C10641, Thermo Fisher Scientific, Inc.) following the manufacturer’s instructions. Following labeling, cells were washed twice with 1× PBS + 0.02% polysorbate 20, then incubated for 2 hours at room temperature in PBS containing 1% goat serum, 2% BSA, 0.05% polysorbate 20, Alexa Fluor^™^ 647-conjugated phalloidin (#A22287, Thermo Fisher Scientific Inc.), and DAPI (#62248, Thermo Fisher Scientific Inc.) to stain F-actin and nuclei, respectively. Final washes included three rinses with PBS + 0.02% polysorbate 20, one rinse with PBS alone, and storage in PBS containing 1X VectCell^™^ Trolox (#CB-1000-2, Vector Laboratories, Inc.) until imaging.

**Microscopy and image quantification.** Confocal images were acquired on a Zeiss LSM 710 confocal microscope using a 60× oil immersion objective and 405, 488, and 647 nm laser lines. For ASO localization and quantification, images were processed using ImageJ software, v1.54^51^, Cellpose software, v2.2.3^95^, ilastik software, v1.4.0^96^, and CellProfiler software, v4.2.6^97^. Raw single-channel images and overlays were first extracted in ImageJ software. Cell segmentation masks were generated using Cellpose software: composite DAPI and phalloidin images were used to train a custom model for cytoplasmic segmentation, while nuclei were segmented using the built-in pretrained nuclei model. To exclude ASO aggregates from quantification, ilastik software was trained on a representative subset of images to identify large fluorescent aggregates, and the resulting ASO aggregation masks were applied to the full dataset. All images and segmentation masks were imported into CellProfiler software for quantification.

Individual cells were identified using Cellpose software-derived masks, and ASO aggregates were excluded based on ilastik software-generated masks. Fluorescent ASO signal was quantified per nucleus across segmented cells. Reported results represent the relative mean nuclear ASO intensity per cell, normalized to the total cellular ASO intensity in non-targeting control (NTC) samples at 24 hours.

Relative intracellular ASO intensities (fold changes relative to untreated control samples) were analyzed to assess differences in temporal expression profiles between treatment groups. A linear mixed-effects model implemented in the nlme package (v3.1-168) in R (v4.2.2) was used to account for the repeated-measures structure of the experiment, where the same biological replicates were measured at multiple time points.

The model included Time, Treatment, and their interaction (Time × Treatment) as fixed effects, with replicate ID specified as a random effect to model intra-subject correlation. The model formula was: Concentration ∼ Time * Treatment, random = ∼1 | Replicate_ID

Model residuals were inspected visually to confirm normality and homoscedasticity. Statistical significance of fixed effects was assessed using ANOVA on the fitted model, with the primary test of interest being the Time × Treatment interaction, which indicates whether the temporal trajectory of gene expression differs between treatments.

All statistical tests were two-sided, and a p-value < 0.05 was considered significant. Data visualization was performed using Prism software version 10.6.0 with expression values plotted as mean FC ± SD at each time point.

**ChIP-seq data processing**. ChIP-seq FASTQ files were processed using the nf-core/chipseq pipeline (v2.0.0). The pipeline was executed on an AWS Batch cloud environment using containers to ensure full reproducibility. Sequencing reads were first quality-checked using FastQC (v0.11.9). Adapter trimming and quality filtering were performed using Trim Galore! (v0.6.7). High-quality reads were aligned to the reference genome (e.g., hg38) using BWA-MEM (v0.7.17). Duplicates were marked and removed using Picard (v2.27.4), and alignment quality metrics were generated via samtools (v1.15.1) and deepTools (v3.5.1). Peaks were called using MACS2 (v2.2.7.1) with appropriate input controls. For histone modifications, broad peaks were called using the default mode, while transcription factors were processed in narrow peak mode by adding the –-narrow_peak label during peak calling. Signal tracks (bigWig) and peak were annotated using HOMER (v4.11)^98^ to support downstream analysis. Input samples were used to normalize background signal. Quality control metrics were aggregated using MultiQC (v1.13). All workflow parameters and software versions were recorded by the pipeline to ensure transparency and reproducibility.

**RNA-seq data processing.** RNA-seq data were processed using the nf-core/rnaseq pipeline (v3.14.0). The pipeline was executed with default parameters unless otherwise specified. Briefly, raw sequencing reads were subjected to quality control and adapter trimming using FastQC (v0.11.9) and Cutadapt (v4.4)^99^, followed by alignment to the reference genome (hg38) using STAR (v2.7.10a). Gene– and transcript-level quantification was performed using RSEM (v1.3.3), and feature-based counts were obtained with featureCounts (v2.0.3) based on the GENCODE v43 annotation. Quality control reports from all steps were aggregated using MultiQC (v1.14). Strand-specific normalized coverage files in bigWig format were generated using deepTools (v3.5.4) to facilitate visualization.

**PRO-seq data processing.** PRO-seq data were processed using the nf-core/nascent pipeline (version 2.2.0),. The pipeline was executed with default parameters unless otherwise specified. Briefly, raw reads were subjected to adapter and quality trimming using fastp (v0.23.4). Trimmed reads were aligned to the reference genome (hg38) using BWA (v0.7.17). Unique molecular identifiers (UMIs) were extracted and UMI-based deduplication was performed using UMI-tools (v1.1.5). Duplicate read marking was carried out with Picard MarkDuplicates (v2.27.5). Strand-specific coverage graphs were generated by creating bedGraph files with BEDTools (v2.30.0) and converting them to bigWig format using deepTools (v3.5.1). Quality control metrics were generated using MultiQC (v1.20), integrating outputs from various steps such as read quality, mapping statistics, and duplication rates.

**ATAC-seq data processing**. ATAC-seq data were processed using the nf-core/atacseq pipeline (v2.1.2). The pipeline was executed in a containerized environment using Docker via AWS Batch to ensure scalability and reproducibility. Paired-end FASTQ reads were first subjected to FastQC (v0.11.9) for initial quality assessment. Adapter sequences and low-quality bases were trimmed using Trim Galore! (v0.6.7). High-quality reads were aligned to the reference genome (e.g., hg38) using BWA-MEM (v0.7.17). PCR duplicates were marked and removed using Picard (v3.0.0), and mitochondrial reads were filtered out to focus the analyses on nuclear chromatin accessibility. Post-alignment quality metrics were computed using samtools (v1.17) and deepTools (v3.5.1). Fragment size distribution, TSS enrichment scores, and other ATAC-seq-specific quality metrics were calculated to evaluate library quality. Peaks were called using MACS2 (v2.2.7.1) with –-narrow_peak label. Signal tracks in bigWig format were generated for visualization, and peaks were annotated using HOMER (v4.11) and BEDTools (v2.30.0). Final quality control metrics were compiled into an interactive report using MultiQC (v1.13).

**Iso-Seq data processing and transcript analysis.** Raw subreads generated from the PacBio Sequel II platform were processed using command-line utilities provided in SMRT Link version 13.0 (Pacific Biosciences of California, Inc.). The analysis followed the Iso-Seq workflow, with additional steps for alignment, transcript curation, and quality control.

**CCS generation:** subreads were first converted into high-fidelity (HiFi) reads using the ccs tool. Reads were required to meet a minimum predicted accuracy threshold of 0.80 and fall within a length range of 50 to 50,000 nts. The setting for minimum number of passes was set to zero, allowing inclusion of low-pass reads for completeness. *[--min-rq 0.80, –-min-passes 0, –-min-length 50, –-max-length 50000, –-all-kinetics, –-hifi-kinetics]*

**Primer trimming:** full-length cDNA reads were identified by removing primer sequences using lima, which was run in Iso-Seq mode. A FASTA file containing primer sequences was used for this step.

**PolyA Tail Removal and FLNC Selection:** next, isoseq refine was applied to trim polyadenylation tails and enrich for full-length non-chimeric (FLNC) reads. Only reads containing detectable polyA sequences of at least 12 bases were retained. This step ensured the inclusion of mature transcript ends. *[--require-polya, –-min-polya-length]*

**Alignment to the reference genome:** cleaned FLNC reads were mapped to the hg38 human genome assembly using pbmm2 with the ISOSEQ preset for optimized performance. Alignments were sorted and summarized with samtools to assess mapping quality. *[-N 1, –-preset ISOSEQ, –-sort]*

**Transcript extraction and annotation:** aligned reads were converted to BED12 format with BEDTools, followed by conversion to GTF using a custom Perl script. This provided an initial annotation of transcript structures based on read alignments.

**Clustering of isoforms:** transcript isoforms were clustered using isoseq cluster2, generating both high-confidence polished transcripts and unclustered singleton reads. These were then realigned to the genome to produce a merged alignment file. *[--singletons]*

**Appending read counts:** a custom R script was used to append read count information to each transcript’s name in the merged alignment file to keep track of enrichment. The updated BAM file was reindexed for downstream compatibility. *[addCountsToBamTracks_v2.R]*

**Isoform collapsing and de-redundancy:** transcript models were then collapsed using isoseq collapse, which groups similar isoforms (clusters + singletons) while preserving differences in transcript boundaries. Parameters were set to allow for slight variations (up to 5 nt) in both 5′ and 3′ ends, without collapsing transcripts with additional upstream exons.

*[--min-aln-coverage=0, –-min-aln-identity=0, –-do-not-collapse-extra-5exons, –-max-5p-diff 5, –-max-3p-diff 5]*

**Conversion to standard formats:** the collapsed transcripts were converted into GTF format using gffread, facilitating compatibility with other genome analysis tools.

**Transcript quality control and annotation with SQANTI3**^98^: to evaluate, annotate and filter the resulting isoforms, the SQANTI3 toolkit (v5.1.1) was employed. Transcript models were annotated and classified based on known GENCODE v43 annotations, and multiple layers of quality metrics were applied, including support from polyA motifs, known polyA peaks, and CAGE data. Filtering was done using custom rule sets, and rescued mono-exonic isoforms were recovered in a final pass. According to SQANTI3 annotation, transcripts classified as Full Splice Match (FSM), Incomplete Splice Match (ISM), Genic, Genic Intron, or Fusion were designated as genic mRNAs. All other transcripts were considered novel. Novel transcripts with 5′ ends overlapping one of the 2,000 probed enhancers were defined as regRNAs transcribed from the corresponding enhancers. For paRNAs, only transcripts transcribed in the antisense orientation relative to their associated mRNAs were retained. SQANTI3 assesses the presence and usage of these canonical motifs (GT-AG, GC-AG, AT-AC) versus non-canonical motifs at donor and acceptor sites within the identified splice junctions.

**Final outputs and visualization:** high-confidence transcript annotations were converted to BED format, sorted, and transformed into BigBed format using bedToBigBed with a custom schema.

**SHAPE-MaP Data Processing**. SHAPE-MaP data were processed using ShapeMapper 2 (v2.3)^100^, a publicly available computational pipeline designed for accurate detection of RNA chemical probing reactivities from next-generation sequencing data. The pipeline performs read alignment, mutation detection, and normalization to generate nucleotide-resolution SHAPE reactivity profiles. Paired-end sequencing reads from reverse transcription of chemically modified, untreated, and denatured control RNA samples were provided to ShapeMapper 2 along with reference RNA sequences in FASTA format. The analysis followed the default three-sample SHAPE-MaP workflow, which includes: (1) read preprocessing and merging using *fastp* and *pear*, (2) alignment to the reference sequence with *bowtie2*, (3) mutation detection by base comparison across the three sample types (+, –, and denatured control), and (4) reactivity calculation based on differential mutation rates normalized to the denatured control. The resulting SHAPE reactivity values were used for downstream RNA structure modeling and reactivity-based filtering.

## Acknowledgements

The authors thank Rachel Meyers for helpful comments on the manuscript. This research was funded by CAMP4 Therapeutics Corporation.

## Author contributions

A.A.S., B.N.A., B.J.M., C.M., J.W., S.M., Y.N., E.C., S.W., I.P., E.C., H.T., K.X., G.W., R.K., M.G., E.L., C.J.J., R.P., and J.C. designed and performed the experiments. A.A.S., A.S., D.F.T., B.N.A., B.J.M., Y.L., C.M., J.W., S.M., J.H., Y.N., A.A., E.C., G.G., I.P., Y.G., E.C., H.T., K.X., G.W., R.K., M.G. and E.L. analyzed the data. A.A.S., B.N.A., B.J.M., C.M., J.W., S.M., Y.L., G.G., D.F.T. wrote the manuscript. A.A.S. and D.B. supervised the research. All authors participated in data interpretation.

## References

1. Henninger, J.E., and Young, R.A. (2024). An RNA-centric view of transcription and genome organization. Mol. Cell 84, 3627–3643. 10.1016/j.molcel.2024.08.021.

2. Field, A., and Adelman, K. (2020). Evaluating Enhancer Function and Transcription. Annu. Rev. Biochem. 89, 213–234. 10.1146/annurev-biochem-011420-095916.

3. Heinz, S., Romanoski, C.E., Benner, C., and Glass, C.K. (2015). The selection and function of cell type-specific enhancers. Nat. Rev. Mol. Cell Biol. 16, 144–154. 10.1038/nrm3949.

4. Gasperini, M., Tome, J., and Shendure, J. (2020). Towards a comprehensive catalogue of validated and target-linked human enhancers. Nat. Rev. Genet. 21, 292–310. 10.1038/s41576-019-0209-0.

5. Kim, S., and Wysocka, J. (2023). Deciphering the multi-scale, quantitative cis-regulatory code. Mol. Cell 83, 373–392. 10.1016/j.molcel.2022.12.032.

6. Fulco, C.P., Nasser, J., Jones, T.R., Munson, G., Bergman, D.T., Subramanian, V., Grossman, S.R., Anyoha, R., Doughty, B.R., Patwardhan, T.A., et al. (2019). Activity-by-contact model of enhancer–promoter regulation from thousands of CRISPR perturbations. Nat. Genet. 51, 1664–1669. 10.1038/s41588-019-0538-0.

7. Schoenfelder, S., and Fraser, P. (2019). Long-range enhancer–promoter contacts in gene expression control. Nat. Rev. Genet. 20, 437–455. 10.1038/s41576-019-0128-0.

8. Sigova, A.A., Abraham, B.J., Ji, X., Molinie, B., Hannett, N.M., Guo, Y.E., Jangi, M., Giallourakis, C.C., Sharp, P.A., and Young, R.A. (2015). Transcription factor trapping by RNA in gene regulatory elements. Science 350, 978–981. 10.1126/science.aad3346.

9. Oksuz, O., Henninger, J.E., Warneford-Thomson, R., Zheng, M.M., Erb, H., Vancura, A., Overholt, K.J., Hawken, S.W., Banani, S.F., Lauman, R., et al. (2023). Transcription factors interact with RNA to regulate genes. Mol. Cell 83, 2449–2463.e13. 10.1016/j.molcel.2023.06.012.

10. Long, Y., Wang, X., Youmans, D.T., and Cech, T.R. (2017). How do lncRNAs regulate transcription? Sci. Adv. 3, eaao2110. 10.1126/sciadv.aao2110.

11. Andersson, R., Gebhard, C., Miguel-Escalada, I., Hoof, I., Bornholdt, J., Boyd, M., Chen, Y., Zhao, X., Schmidl, C., Suzuki, T., et al. (2014). An atlas of active enhancers across human cell types and tissues. Nature 507, 455–461. 10.1038/nature12787.

12. Flynn, R.A., Almada, A.E., Zamudio, J.R., and Sharp, P.A. (2011). Antisense RNA polymerase II divergent transcripts are P-TEFb dependent and substrates for the RNA exosome. Proc. Natl. Acad. Sci. 108, 10460–10465. 10.1073/pnas.1106630108.

13. Schaukowitch, K., Joo, J.-Y., Liu, X., Watts, J.K., Martinez, C., and Kim, T.-K. (2014). Enhancer RNA Facilitates NELF Release from Immediate Early Genes. Mol. Cell 56, 29–42. 10.1016/j.molcel.2014.08.023.

14. De Santa, F., Barozzi, I., Mietton, F., Ghisletti, S., Polletti, S., Tusi, B.K., Muller, H., Ragoussis, J., Wei, C.-L., and Natoli, G. (2010). A Large Fraction of Extragenic RNA Pol II Transcription Sites Overlap Enhancers. PLoS Biol. 8, e1000384. 10.1371/journal.pbio.1000384.

15. Schwalb, B., Michel, M., Zacher, B., Frühauf, K., Demel, C., Tresch, A., Gagneur, J., and Cramer, P. (2016). TT-seq maps the human transient transcriptome. Science 352, 1225–1228. 10.1126/science.aad9841.

16. Blumberg, A., Zhao, Y., Huang, Y.-F., Dukler, N., Rice, E.J., Chivu, A.G., Krumholz, K., Danko, C.G., and Siepel, A. (2021). Characterizing RNA stability genome-wide through combined analysis of PRO-seq and RNA-seq data. BMC Biol. 19, 30. 10.1186/s12915-021-00949-x.

17. Mattick, J.S., Amaral, P.P., Carninci, P., Carpenter, S., Chang, H.Y., Chen, L.-L., Chen, R., Dean, C., Dinger, M.E., Fitzgerald, K.A., et al. (2023). Long non-coding RNAs: definitions, functions, challenges and recommendations. Nat. Rev. Mol. Cell Biol. 24, 430–447. 10.1038/s41580-022-00566-8.

18. Clark, M.B., Johnston, R.L., Inostroza-Ponta, M., Fox, A.H., Fortini, E., Moscato, P., Dinger, M.E., and Mattick, J.S. (2012). Genome-wide analysis of long noncoding RNA stability. Genome Res. 22, 885–898. 10.1101/gr.131037.111.

19. Yao, L., Liang, J., Ozer, A., Leung, A.K.-Y., Lis, J.T., and Yu, H. (2022). A comparison of experimental assays and analytical methods for genome-wide identification of active enhancers. Nat. Biotechnol. 40, 1056–1065. 10.1038/s41587-022-01211-7.

20. Duttke, S.H., Chang, M.W., Heinz, S., and Benner, C. (2019). Identification and dynamic quantification of regulatory elements using total RNA. Genome Res. 29, 1836–1846. 10.1101/gr.253492.119.

21. Kopp, F., and Mendell, J.T. (2018). Functional Classification and Experimental Dissection of Long Noncoding RNAs. Cell 172, 393–407. 10.1016/j.cell.2018.01.011.

22. Modulating the expression of long non-coding RNAs for functional studies 10.15252/embr.201846955.

23. Lennox, K.A., and Behlke, M.A. (2016). Cellular localization of long non-coding RNAs affects silencing by RNAi more than by antisense oligonucleotides. Nucleic Acids Res. 44, 863–877. 10.1093/nar/gkv1206.

24. Lam, M.T.Y., Li, W., Rosenfeld, M.G., and Glass, C.K. (2014). Enhancer RNAs and regulated transcriptional programs. Trends Biochem. Sci. 39, 170–182. 10.1016/j.tibs.2014.02.007.

25. Carullo, N.V.N., Phillips III, R.A., Simon, R.C., Soto, S.A.R., Hinds, J.E., Salisbury, A.J., Revanna, J.S., Bunner, K.D., Ianov, L., Sultan, F.A., et al. (2020). Enhancer RNAs predict enhancer–gene regulatory links and are critical for enhancer function in neuronal systems. Nucleic Acids Res. 48, 9550–9570. 10.1093/nar/gkaa671.

26. Statello, L., Guo, C.-J., Chen, L.-L., and Huarte, M. (2021). Gene regulation by long non-coding RNAs and its biological functions. Nat. Rev. Mol. Cell Biol. 22, 96–118. 10.1038/s41580-020-00315-9.

27. Li, W., Notani, D., Ma, Q., Tanasa, B., Nunez, E., Chen, A.Y., Merkurjev, D., Zhang, J., Ohgi, K., Song, X., et al. (2013). Functional roles of enhancer RNAs for oestrogen-dependent transcriptional activation. Nature 498, 516–520. 10.1038/nature12210.

28. Cheng, J.-H., Pan, D.Z.-C., Tsai, Z.T.-Y., and Tsai, H.-K. (2015). Genome-wide analysis of enhancer RNA in gene regulation across 12 mouse tissues. Sci. Rep. 5, 12648. 10.1038/srep12648.

29. Hnisz, D., Abraham, B.J., Lee, T.I., Lau, A., Saint-André, V., Sigova, A.A., Hoke, H., and Young, R.A. (2013). Transcriptional super-enhancers connected to cell identity and disease. Cell 155, 10.1016/j.cell.2013.09.053. https://doi.org/10.1016/j.cell.2013.09.053.

30. Evans, E.F., Shyr, Z.A., Traynor, B.J., and Zheng, W. (2024). Therapeutic development approaches to treat haploinsufficiency diseases: restoring protein levels. Drug Discov. Today 29, 104201. 10.1016/j.drudis.2024.104201.

31. Kim, T.-K., Hemberg, M., and Gray, J.M. (2015). Enhancer RNAs: A Class of Long Noncoding RNAs Synthesized at Enhancers. Cold Spring Harb. Perspect. Biol. 7, a018622. 10.1101/cshperspect.a018622.

32. Lagarde, J., Uszczynska-Ratajczak, B., Carbonell, S., Pérez-Lluch, S., Abad, A., Davis, C., Gingeras, T.R., Frankish, A., Harrow, J., Guigo, R., et al. (2017). High-throughput annotation of full-length long noncoding RNAs with Capture Long-Read Sequencing. Nat. Genet. 49, 1731–1740. 10.1038/ng.3988.

33. Wang, F., Xu, Y., Wang, R., Zhang, B., Smith, N., Notaro, A., Gaerlan, S., Kutschera, E., Kadash-Edmondson, K.E., Xing, Y., et al. (2023). TEQUILA-seq: a versatile and low-cost method for targeted long-read RNA sequencing. Nat. Commun. 14, 4760. 10.1038/s41467-023-40083-6.

34. Iyer, S.V., Goodwin, S., and McCombie, W.R. (2024). Leveraging the power of long reads for targeted sequencing. Genome Res. 34, 1701–1718. 10.1101/gr.279168.124.

35. Henriques, T., Scruggs, B.S., Inouye, M.O., Muse, G.W., Williams, L.H., Burkholder, A.B., Lavender, C.A., Fargo, D.C., and Adelman, K. (2018). Widespread transcriptional pausing and elongation control at enhancers. Genes Dev. 32, 26–41. 10.1101/gad.309351.117.

36. Krivitzky, L., Babikian, T., Lee, H.-S., Thomas, N.H., Burk-Paull, K.L., and Batshaw, M.L. Intellectual, Adaptive, and Behavioral Functioning in Children With Urea Cycle Disorders.

37. Jang, Y.J., LaBella, A.L., Feeney, T.P., Braverman, N., Tuchman, M., Morizono, H., Ah Mew, N., and Caldovic, L. (2018). Disease-causing Mutations in the Promoter and Enhancer of the Ornithine Transcarbamylase Gene. Hum. Mutat. 39, 527–536. 10.1002/humu.23394.

38. Tafessu, A., and Banaszynski, L.A. (2020). Establishment and function of chromatin modification at enhancers. Open Biol. 10, 200255. 10.1098/rsob.200255.

39. Takiguchi, M., and Mori, M. (1995). Transcriptional regulation of genes for ornithine cycle enzymes. Biochem. J. 312, 649–659. 10.1042/bj3120649.

40. Cheng, A.W., Wang, H., Yang, H., Shi, L., Katz, Y., Theunissen, T.W., Rangarajan, S., Shivalila, C.S., Dadon, D.B., and Jaenisch, R. (2013). Multiplexed activation of endogenous genes by CRISPR-on, an RNA-guided transcriptional activator system. Cell Res. 23, 1163–1171. 10.1038/cr.2013.122.

41. Gilbert, L.A., Larson, M.H., Morsut, L., Liu, Z., Brar, G.A., Torres, S.E., Stern-Ginossar, N., Brandman, O., Whitehead, E.H., Doudna, J.A., et al. (2013). CRISPR-Mediated Modular RNA-Guided Regulation of Transcription in Eukaryotes. Cell 154, 442–451. 10.1016/j.cell.2013.06.044.

42. Chardon, F.M., McDiarmid, T.A., Page, N.F., Daza, R.M., Martin, B.K., Domcke, S., Regalado, S.G., Lalanne, J.-B., Calderon, D., Li, X., et al. (2024). Multiplex, single-cell CRISPRa screening for cell type specific regulatory elements. Nat. Commun. 15, 8209. 10.1038/s41467-024-52490-4.

43. Kang, Y.-J., Yang, D.-C., Kong, L., Hou, M., Meng, Y.-Q., Wei, L., and Gao, G. (2017). CPC2: a fast and accurate coding potential calculator based on sequence intrinsic features. Nucleic Acids Res. 45, W12–W16. 10.1093/nar/gkx428.

44. Viollet, B., Foretz, M., Guigas, B., Horman, S., Dentin, R., Bertrand, L., Hue, L., and Andreelli, F. (2006). Activation of AMP-activated protein kinase in the liver: a new strategy for the management of metabolic hepatic disorders. J. Physiol. 574, 41–53. 10.1113/jphysiol.2006.108506.

45. Heibel, S.K., McGuire, P.J., Haskins, N., Datta Majumdar, H., Rayavarapu, S., Nagaraju, K., Hathout, Y., Brown, K., Tuchman, M., and Caldovic, L. (2019). AMPK Signaling Regulates Expression of Urea Cycle Enzymes in Response to Changes in Dietary Protein Intake. J. Inherit. Metab. Dis. 42, 1088–1096. 10.1002/jimd.12133.

46. Ahn, S.H., Lee, Y.-J., Lim, D.S., Cho, W., Gwon, H.J., Abd El-Aty, A.M., Jeong, J.H., and Jung, T.W. (2024). Upadacitinib counteracts hepatic lipid deposition *via* the repression of JAK1/STAT3 signaling and AMPK/autophagy-mediated suppression of ER stress. Biochem. Biophys. Res. Commun. 735, 150829. 10.1016/j.bbrc.2024.150829.

47. Liang, X.-H., Sun, H., Nichols, J.G., and Crooke, S.T. (2017). RNase H1-Dependent Antisense Oligonucleotides Are Robustly Active in Directing RNA Cleavage in Both the Cytoplasm and the Nucleus. Mol. Ther. 25, 2075–2092. 10.1016/j.ymthe.2017.06.002.

48. Takeshima, Y., Nishio, H., Sakamoto, H., Nakamura, H., and Matsuo, M. (1995). Modulation of in vitro splicing of the upstream intron by modifying an intra-exon sequence which is deleted from the dystrophin gene in dystrophin Kobe. J. Clin. Invest. 95, 515–520. 10.1172/jci117693.

49. Shen, X., and Corey, D.R. (2018). Chemistry, mechanism and clinical status of antisense oligonucleotides and duplex RNAs. Nucleic Acids Res. 46, 1584–1600. 10.1093/nar/gkx1239.

50. Sang, A., Zhuo, S., Bochanis, A., Manautou, J.E., Bahal, R., Zhong, X., and Rasmussen, T.P. (2024). Mechanisms of Action of the US Food and Drug Administration-Approved Antisense Oligonucleotide Drugs. BioDrugs Clin. Immunother. Biopharm. Gene Ther. 38, 511–526. 10.1007/s40259-024-00665-2.

51. Roberts, T.C., Langer, R., and Wood, M.J.A. (2020). Advances in oligonucleotide drug delivery. Nat. Rev. Drug Discov. 19, 673–694. 10.1038/s41573-020-0075-7.

52. Micheletti, R., Plaisance, I., Abraham, B.J., Sarre, A., Ting, C.-C., Alexanian, M., Maric, D., Maison, D., Nemir, M., Young, R.A., et al. (2017). The long noncoding RNA Wisper controls cardiac fibrosis and remodeling. Sci. Transl. Med. 9, eaai9118. 10.1126/scitranslmed.aai9118.

53. Stein, C.A., Hansen, J.B., Lai, J., Wu, S., Voskresenskiy, A., H⊘g, A., Worm, J., Hedtjärn, M., Souleimanian, N., Miller, P., et al. (2010). Efficient gene silencing by delivery of locked nucleic acid antisense oligonucleotides, unassisted by transfection reagents. Nucleic Acids Res. 38, e3–e3. 10.1093/nar/gkp841.

54. Partridge, E.C., Chhetri, S.B., Prokop, J.W., Ramaker, R.C., Jansen, C.S., Goh, S.-T., Mackiewicz, M., Newberry, K.M., Brandsmeier, L.A., Meadows, S.K., et al. (2020). Occupancy maps of 208 chromatin-associated proteins in one human cell type. Nature 583, 720–728. 10.1038/s41586-020-2023-4.

55. Raab, J.R., Resnick, S., and Magnuson, T. (2015). Genome-Wide Transcriptional Regulation Mediated by Biochemically Distinct SWI/SNF Complexes. PLoS Genet. 11, e1005748. 10.1371/journal.pgen.1005748.

56. Doetzlhofer, A., Rotheneder, H., Lagger, G., Koranda, M., Kurtev, V., Brosch, G., Wintersberger, E., and Seiser, C. (1999). Histone deacetylase 1 can repress transcription by binding to Sp1. Mol. Cell. Biol. 19, 5504–5511. 10.1128/MCB.19.8.5504.

57. O’Connor, L., Gilmour, J., and Bonifer, C. (2016). The Role of the Ubiquitously Expressed Transcription Factor Sp1 in Tissue-specific Transcriptional Regulation and in Disease. Yale J. Biol. Med. 89, 513–525.

58. Ci, W., Polo, J.M., Cerchietti, L., Shaknovich, R., Wang, L., Yang, S.N., Ye, K., Farinha, P., Horsman, D.E., Gascoyne, R.D., et al. (2009). The BCL6 transcriptional program features repression of multiple oncogenes in primary B cells and is deregulated in DLBCL. Blood 113, 5536–5548. 10.1182/blood-2008-12-193037.

59. Odnokoz, O., Wavelet-Vermuse, C., Hophan, S.L., Bulun, S., and Wan, Y. ARID1 proteins: from transcriptional and post-translational regulation to carcinogenesis and potential therapeutics. Epigenomics 13, 809–823. 10.2217/epi-2020-0414.

60. Kelly, R.D.W., Chandru, A., Watson, P.J., Song, Y., Blades, M., Robertson, N.S., Jamieson, A.G., Schwabe, J.W.R., and Cowley, S.M. (2018). Histone deacetylase (HDAC) 1 and 2 complexes regulate both histone acetylation and crotonylation in vivo. Sci. Rep. 8, 14690. 10.1038/s41598-018-32927-9.

61. Wang, C., Guo, Z., Chu, C., Lu, Y., Zhang, X., and Zhan, X. (2023). Two assembly modes for SIN3 histone deacetylase complexes. Cell Discov. 9, 42. 10.1038/s41421-023-00539-x.

62. Basta, J., and Rauchman, M. (2015). The Nucleosome Remodeling and Deacetylase (NuRD) Complex in Development and Disease. Transl. Res. J. Lab. Clin. Med. 165, 36–47. 10.1016/j.trsl.2014.05.003.

63. Maksour, S., Ooi, L., and Dottori, M. (2020). More than a Corepressor: The Role of CoREST Proteins in Neurodevelopment. eNeuro 7, ENEURO.0337-19.2020. 10.1523/ENEURO.0337-19.2020.

64. Mondal, B., Jin, H., Kallappagoudar, S., Sedkov, Y., Martinez, T., Sentmanat, M.F., Poet, G.J., Li, C., Fan, Y., Pruett-Miller, S.M., et al. The histone deacetylase complex MiDAC regulates a neurodevelopmental gene expression program to control neurite outgrowth. eLife 9, e57519. 10.7554/eLife.57519.

65. Grozinger, C.M., and Schreiber, S.L. (2000). Regulation of histone deacetylase 4 and 5 and transcriptional activity by 14-3-3-dependent cellular localization. Proc. Natl. Acad. Sci. 97, 7835–7840. 10.1073/pnas.140199597.

66. Mottis, A., Mouchiroud, L., and Auwerx, J. (2013). Emerging roles of the corepressors NCoR1 and SMRT in homeostasis. Genes Dev. 27, 819–835. 10.1101/gad.214023.113.

67. Paluvai, H., Shanmukha, K.D., Tyedmers, J., and Backs, J. (2023). Insights into the function of HDAC3 and NCoR1/NCoR2 co-repressor complex in metabolic diseases. Front. Mol. Biosci. 10, 1190094. 10.3389/fmolb.2023.1190094.

68. Ramboer, E., Vanhaecke, T., Rogiers, V., and Vinken, M. (2015). Immortalized human hepatic cell lines for in vitro testing and research purposes. Methods Mol. Biol. Clifton NJ 1250, 53–76. 10.1007/978-1-4939-2074-7_4.

69. Core, L.J., Martins, A.L., Danko, C.G., Waters, C., Siepel, A., and Lis, J.T. (2014). Analysis of nascent RNA identifies a unified architecture of initiation regions at mammalian promoters and enhancers. Nat. Genet. 46, 1311–1320. 10.1038/ng.3142.

70. Andersson, R., Sandelin, A., and Danko, C.G. (2015). A unified architecture of transcriptional regulatory elements. Trends Genet. 31, 426–433. 10.1016/j.tig.2015.05.007.

71. Li, W., Notani, D., and Rosenfeld, M.G. (2016). Enhancers as non-coding RNA transcription units: recent insights and future perspectives. Nat. Rev. Genet. 17, 207–223. 10.1038/nrg.2016.4.

72. Mikhaylichenko, O., Bondarenko, V., Harnett, D., Schor, I.E., Males, M., Viales, R.R., and Furlong, E.E.M. (2018). The degree of enhancer or promoter activity is reflected by the levels and directionality of eRNA transcription. Genes Dev. 32, 42–57. 10.1101/gad.308619.117.

73. Mousavi, K., Zare, H., Dell’Orso, S., Grontved, L., Gutierrez-Cruz, G., Derfoul, A., Hager, G.L., and Sartorelli, V. (2013). eRNAs Promote Transcription by Establishing Chromatin Accessibility at Defined Genomic Loci. Mol. Cell 51, 606–617. 10.1016/j.molcel.2013.07.022.

74. So, B.R., Di, C., Cai, Z., Venters, C.C., Guo, J., Oh, J.-M., Arai, C., and Dreyfuss, G. (2019). A Complex of U1 snRNP with Cleavage and Polyadenylation Factors Controls Telescripting, Regulating mRNA Transcription in Human Cells. Mol. Cell 76, 590–599.e4. 10.1016/j.molcel.2019.08.007.

75. Wu, G., Schmid, M., Rib, L., Polak, P., Meola, N., Sandelin, A., and Jensen, T.H. (2020). A Two-Layered Targeting Mechanism Underlies Nuclear RNA Sorting by the Human Exosome. Cell Rep. 30, 2387–2401.e5. 10.1016/j.celrep.2020.01.068.

76. Melé, M., Mattioli, K., Mallard, W., Shechner, D.M., Gerhardinger, C., and Rinn, J.L. (2017). Chromatin environment, transcriptional regulation, and splicing distinguish lincRNAs and mRNAs. Genome Res. 27, 27–37. 10.1101/gr.214205.116.

77. Basu, K., Dey, A., and Kiran, M. (2023). Inefficient splicing of long non-coding RNAs is associated with higher transcript complexity in human and mouse. RNA Biol. 20, 563–572. 10.1080/15476286.2023.2242649.

78. Kim, T.-K., Hemberg, M., Gray, J.M., Costa, A.M., Bear, D.M., Wu, J., Harmin, D.A., Laptewicz, M., Barbara-Haley, K., Kuersten, S., et al. (2010). Widespread transcription at neuronal activity-regulated enhancers. Nature 465, 182–187. 10.1038/nature09033.

79. Bennett, C.F., and Swayze, E.E. (2010). RNA Targeting Therapeutics: Molecular Mechanisms of Antisense Oligonucleotides as a Therapeutic Platform. Annu. Rev. Pharmacol. Toxicol. 50, 259–293. 10.1146/annurev.pharmtox.010909.105654.

80. Wahlestedt, C. (2013). Targeting long non-coding RNA to therapeutically upregulate gene expression. Nat. Rev. Drug Discov. 12, 433–446. 10.1038/nrd4018.

81. Rinaldi, C., and Wood, M.J.A. (2018). Antisense oligonucleotides: the next frontier for treatment of neurological disorders. Nat. Rev. Neurol. 14, 9–21. 10.1038/nrneurol.2017.148.

82. Crooke, S.T., Baker, B.F., Crooke, R.M., and Liang, X. (2021). Antisense technology: an overview and prospectus. Nat. Rev. Drug Discov. 20, 427–453. 10.1038/s41573-021-00162-z.

83. Crooke, S.T., Liang, X.-H., Baker, B.F., and Crooke, R.M. (2021). Antisense technology: A review. J. Biol. Chem. 296. 10.1016/j.jbc.2021.100416.

84. Lim, K.H., Han, Z., Jeon, H.Y., Kach, J., Jing, E., Weyn-Vanhentenryck, S., Downs, M., Corrionero, A., Oh, R., Scharner, J., et al. (2020). Antisense oligonucleotide modulation of non-productive alternative splicing upregulates gene expression. Nat. Commun. 11, 3501. 10.1038/s41467-020-17093-9.

85. Hill, B., Jaques, M.R., Nair, R.R., Whiffin, N., Wood, M.J.A., Sanders, S.J., Oliver, P.L., Hill, A.C., Rinaldi, C., and Consortium, the U. (2025). Accurately modelling RNase H-mediated antisense oligonucleotide efficacy. Preprint at bioRxiv, 10.1101/2025.10.29.685292 https://doi.org/10.1101/2025.10.29.685292.

86. Bennett, C.F., Baker, B.F., Pham, N., Swayze, E., and Geary, R.S. (2017). Pharmacology of Antisense Drugs. Annu. Rev. Pharmacol. Toxicol. 57, 81–105. 10.1146/annurev-pharmtox-010716-104846.

87. Gorbovytska, V., Kim, S.-K., Kuybu, F., Götze, M., Um, D., Kang, K., Pittroff, A., Brennecke, T., Schneider, L.-M., Leitner, A., et al. (2022). Enhancer RNAs stimulate Pol II pause release by harnessing multivalent interactions to NELF. Nat. Commun. 13, 2429. 10.1038/s41467-022-29934-w.

88. Kole, R., Krainer, A.R., and Altman, S. (2012). RNA therapeutics: Beyond RNA interference and antisense oligonucleotides. Nat. Rev. Drug Discov. 11, 125–140. 10.1038/nrd3625.

89. Havens, M.A., and Hastings, M.L. (2016). Splice-switching antisense oligonucleotides as therapeutic drugs. Nucleic Acids Res. 44, 6549–6563. 10.1093/nar/gkw533.

90. Datlinger, P., Rendeiro, A.F., Schmidl, C., Krausgruber, T., Traxler, P., Klughammer, J., Schuster, L.C., Kuchler, A., Alpar, D., and Bock, C. (2017). Pooled CRISPR screening with single-cell transcriptome readout. Nat. Methods 14, 297–301. 10.1038/nmeth.4177.

91. Livak, K.J., and Schmittgen, T.D. (2001). Analysis of relative gene expression data using real-time quantitative PCR and the 2(-Delta Delta C(T)) Method. Methods San Diego Calif 25, 402–408. 10.1006/meth.2001.1262.

92. Li, X., Yu, K., Li, F., Lu, W., Wang, Y., Zhang, W., and Bai, Y. (2022). Novel Method of Full-Length RNA-seq That Expands the Identification of Non-Polyadenylated RNAs Using Nanopore Sequencing. Anal. Chem. 94, 12342–12351. 10.1021/acs.analchem.2c01128.

93. Buenrostro, J., Wu, B., Chang, H., and Greenleaf, W. (2015). ATAC-seq: A Method for Assaying Chromatin Accessibility Genome-Wide. Curr. Protoc. Mol. Biol. Ed. Frederick M Ausubel Al 109, 21.29.1–21.29.9. 10.1002/0471142727.mb2129s109.

94. Mimoso, C.A., and Goldman, S.R. (2023). PRO-seq: Precise Mapping of Engaged RNA Pol II at Single-Nucleotide Resolution. Curr. Protoc. 3, e961. 10.1002/cpz1.961.

95. Stringer, C., Wang, T., Michaelos, M., and Pachitariu, M. (2021). Cellpose: a generalist algorithm for cellular segmentation. Nat. Methods 18, 100–106. 10.1038/s41592-020-01018-x.

96. Berg, S., Kutra, D., Kroeger, T., Straehle, C.N., Kausler, B.X., Haubold, C., Schiegg, M., Ales, J., Beier, T., Rudy, M., et al. (2019). ilastik: interactive machine learning for (bio)image analysis. Nat. Methods 16, 1226–1232. 10.1038/s41592-019-0582-9.

97. Stirling, D.R., Swain-Bowden, M.J., Lucas, A.M., Carpenter, A.E., Cimini, B.A., and Goodman, A. (2021). CellProfiler 4: improvements in speed, utility and usability. BMC Bioinformatics 22, 433. 10.1186/s12859-021-04344-9.

98. Heinz, S., Benner, C., Spann, N., Bertolino, E., Lin, Y.C., Laslo, P., Cheng, J.X., Murre, C., Singh, H., and Glass, C.K. (2010). Simple Combinations of Lineage-Determining Transcription Factors Prime cis-Regulatory Elements Required for Macrophage and B Cell Identities. Mol. Cell 38, 576–589. 10.1016/j.molcel.2010.05.004.

99. Martin, M. (2011). Cutadapt removes adapter sequences from high-throughput sequencing reads. EMBnet.journal 17, 10–12. 10.14806/ej.17.1.200.

100. Busan, S., and Weeks, K.M. (2018). Accurate detection of chemical modifications in RNA by mutational profiling (MaP) with ShapeMapper 2. RNA N. Y. N 24, 143–148. 10.1261/rna.061945.117.

