## Supplemental Figures for "Profiling and Targeting of Regulatory RNAs to Upregulate Gene Expression"

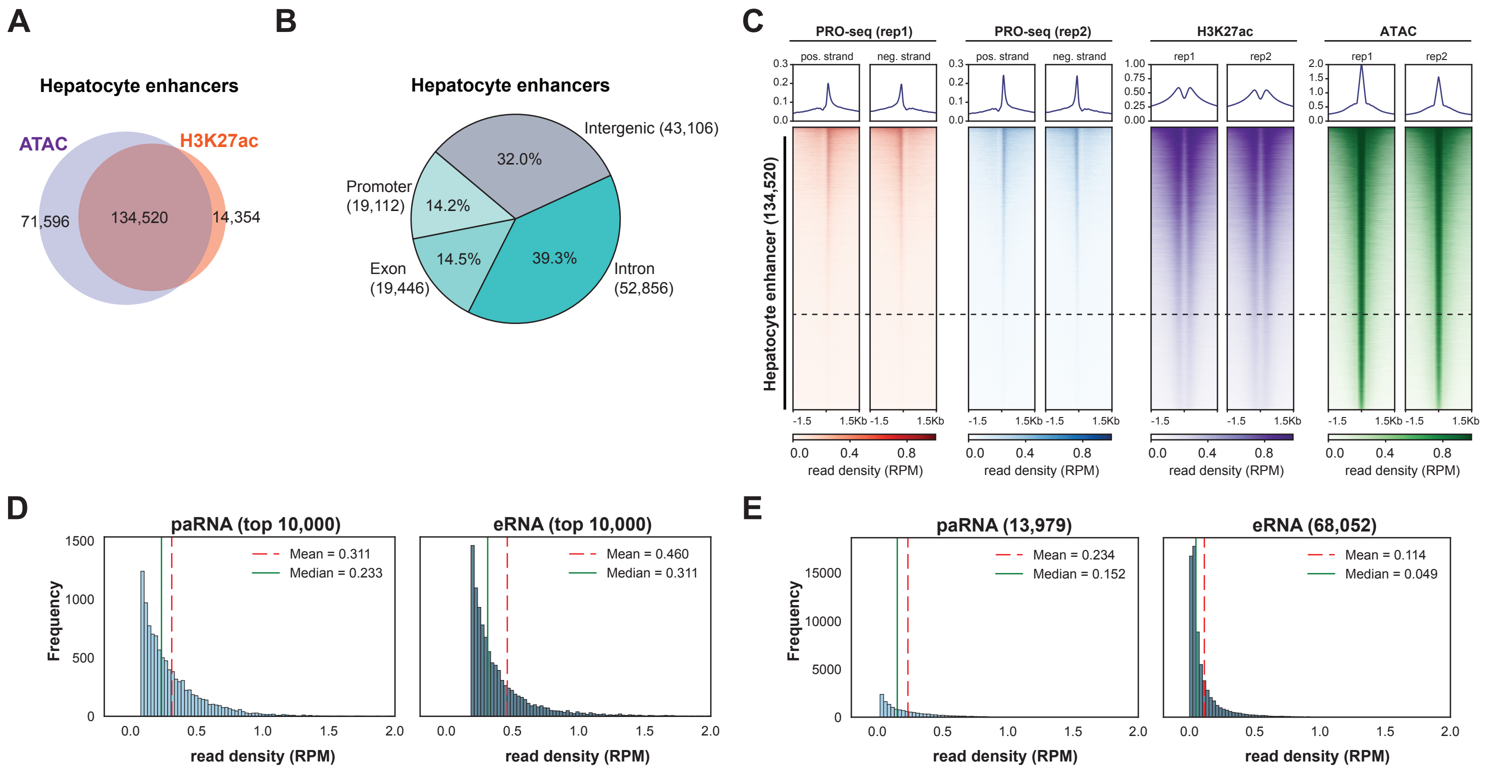


Supplementary Figure 1

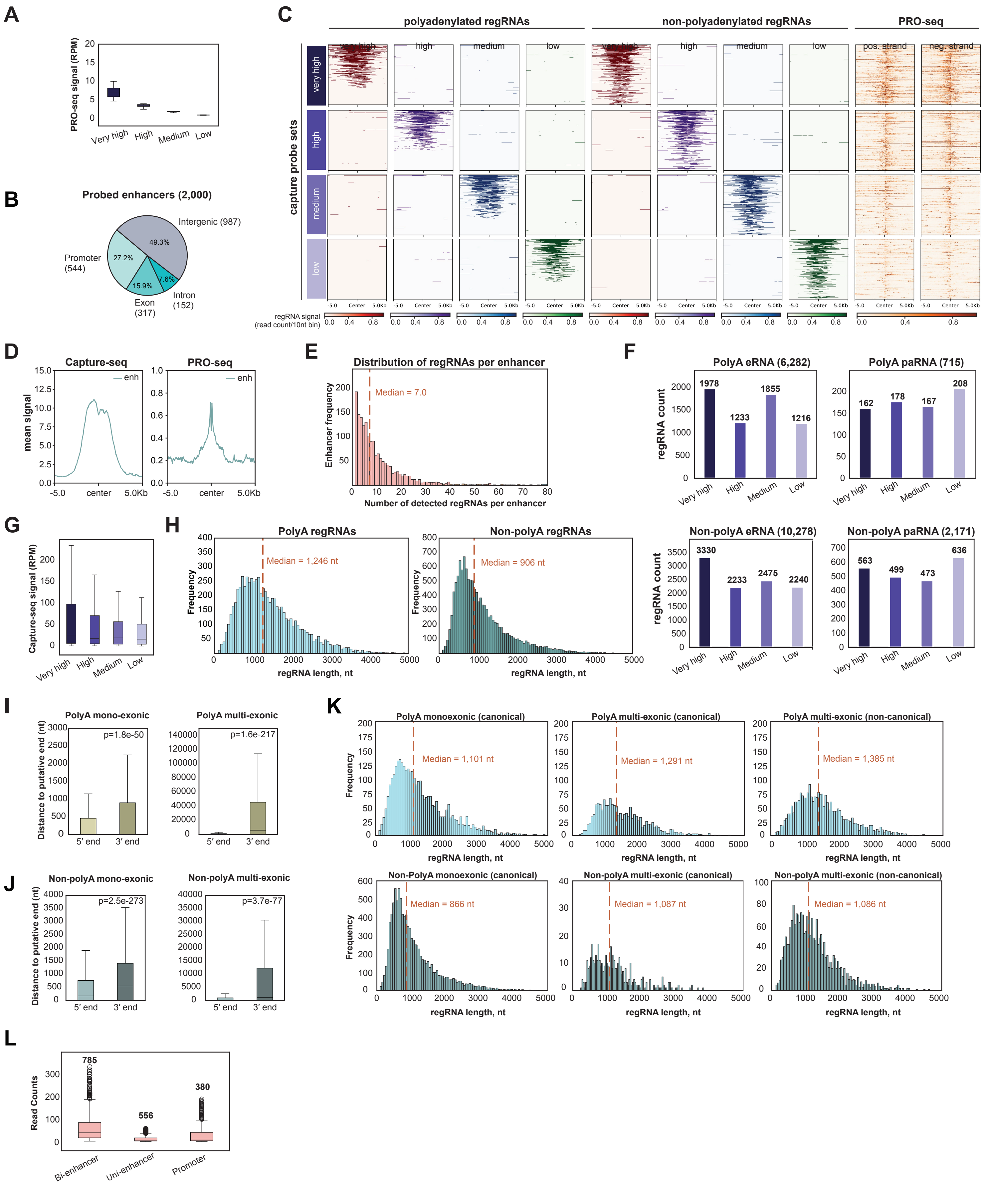


Supplementary Figure 2

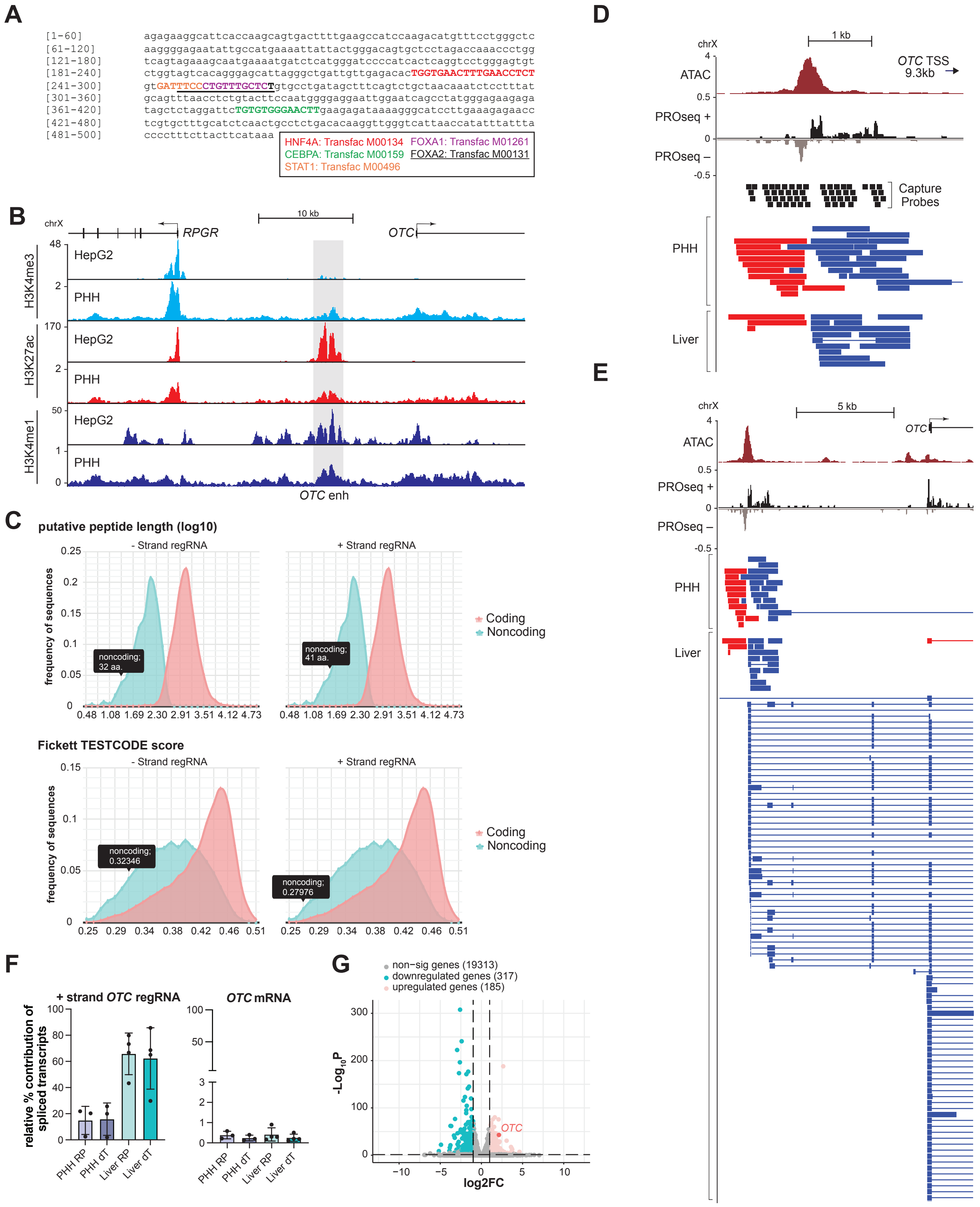


Supplementary Figure 3

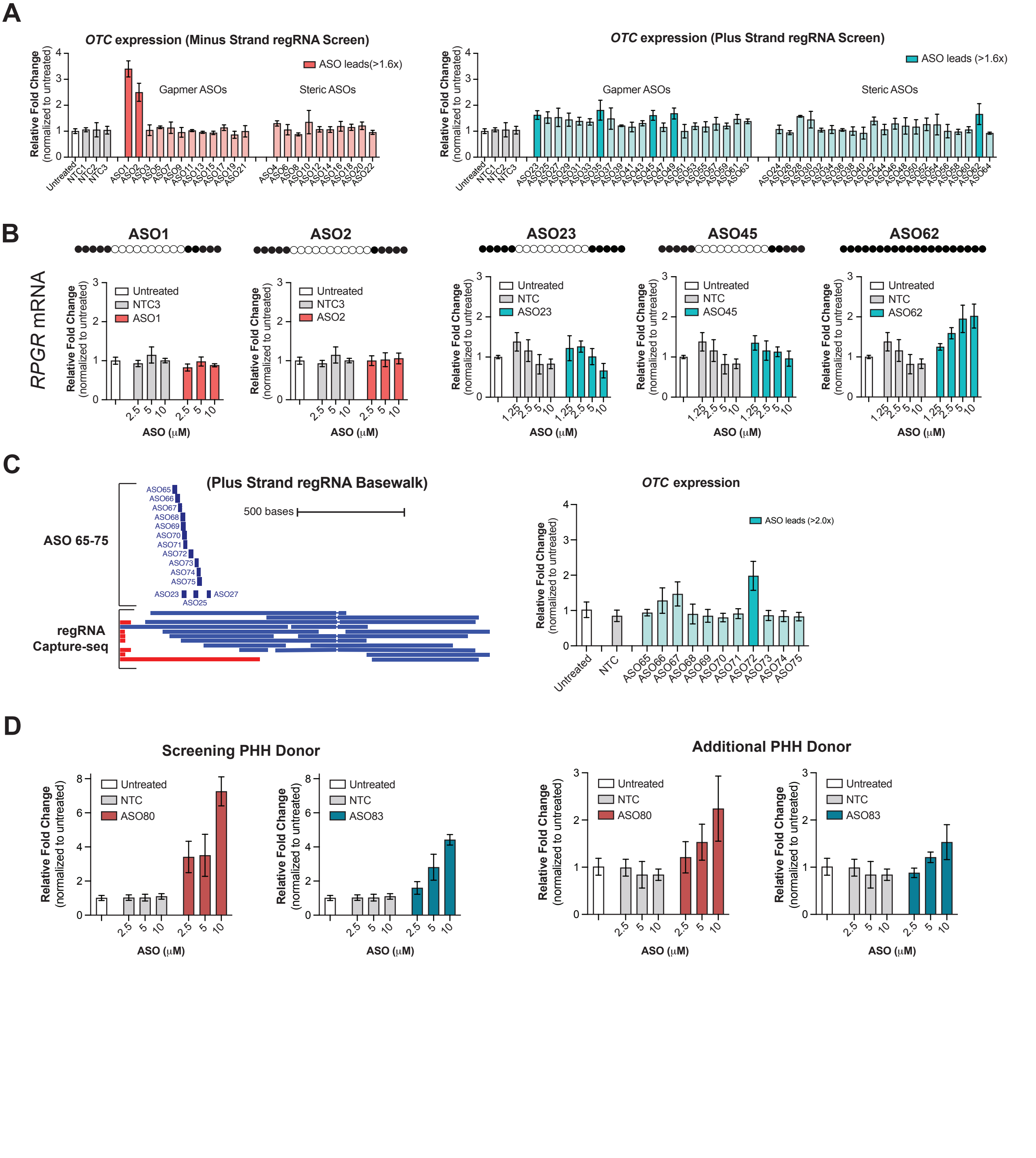


Supplementary Figure 4

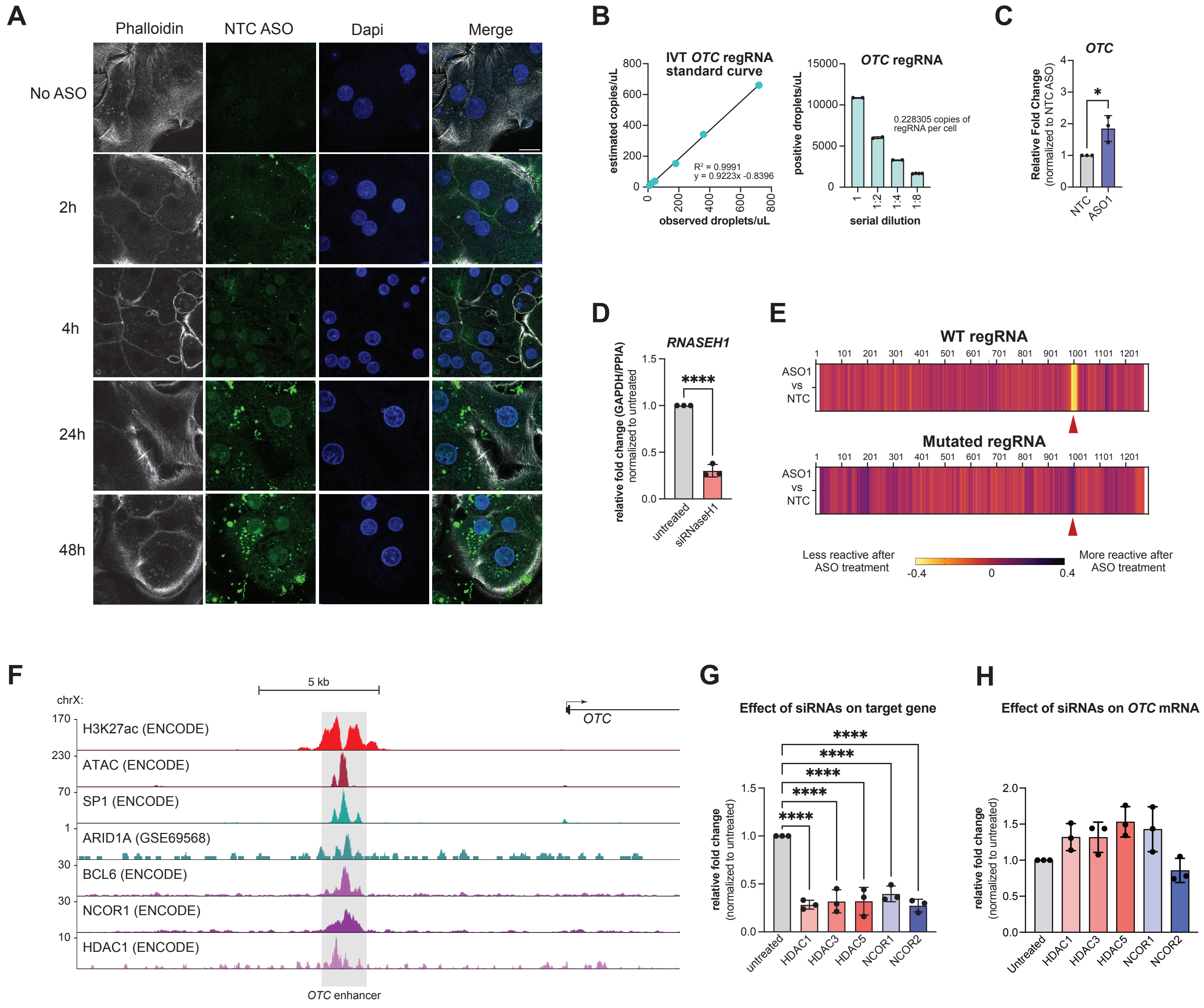


Supplementary Figure 5
