## Supplemental Tables for "Profiling and Targeting of Regulatory RNAs to Upregulate Gene Expression"

Supplementary Table 1

| **gRNA** | **Sequence** |
|---|---|
| *OTC* TSS | 5´-CCTGCAGTATCTCTAACCAG-3’ |
| *OTC* enh | 5´-GATCCCCATCACTCAGGTCC-3’ |
| NTC | 5´-GTTCCGCGTTACATAACTTA-3´ |

**Supplementary Table 2: Commercial TaqMan Probes for qPCR**

| **Target** | **Catalog number** |
|---|---|
| *B2M* | 4326319E |
| *GAPDH* | 4326317E |
| *HDAC1* | Hs02621185_m1 |
| *HDAC3* | Hs00187320_m1 |
| *HDAC5* | Hs00608351_m1 |
| *NCOR1* | Hs01094541_m1 |
| *NCOR2* | Hs00196955_m1 |
| *OTC* | Hs00166892_m1 |
| *PPIA* | 4326316E |
| *RNaseH1* | Hs00268000_m1 |

Supplementary Table 3: ASO sequences

| ASO ID | Sequence | Design | Modifications |
|---|---|---|---|
| ASO1 | TTAATACAGCTCTGGAGTGG | Gapmer | MMMMMDDDDDDDDDDMMMMM |
| ASO2 | AATACAGCTCTGGAGTGGGGT | Gapmer | MMMMMDDDDDDDDDDDMMMMM |
| ASO3 | CGGACACCTCAACACTTTTA | Gapmer | MMMMMDDDDDDDDDDMMMMM |
| ASO4 | CGGACACCTCAACACTTTTA | Steric | MMMMMMMMMMMMMMMMMMMM |
| ASO5 | GGTAGTAGTTAACAAAAGCT | Gapmer | MMMMMDDDDDDDDDDMMMMM |
| ASO6 | GGTAGTAGTTAACAAAAGCT | Steric | MMMMMMMMMMMMMMMMMMMM |
| ASO7 | CTACAGTACTCTCTATTCAT | Gapmer | MMMMMDDDDDDDDDDMMMMM |
| ASO8 | CTACAGTACTCTCTATTCAT | Steric | MMMMMMMMMMMMMMMMMMMM |
| ASO9 | CGCTTACTTCTTAATGGTAA | Gapmer | MMMMMDDDDDDDDDDMMMMM |
| ASO10 | CGCTTACTTCTTAATGGTAA | Steric | MMMMMMMMMMMMMMMMMMMM |
| ASO11 | GCATAACAATGAAGGTGACC | Gapmer | MMMMMDDDDDDDDDDMMMMM |
| ASO12 | GCATAACAATGAAGGTGACC | Steric | MMMMMMMMMMMMMMMMMMMM |
| ASO13 | AGGATTCCCATGGTCTATCT | Gapmer | MMMMMDDDDDDDDDDMMMMM |
| ASO14 | AGGATTCCCATGGTCTATCT | Steric | MMMMMMMMMMMMMMMMMMMM |
| ASO15 | ATAGGTCCCATCTTTACAGG | Gapmer | MMMMMDDDDDDDDDDMMMMM |
| ASO16 | ATAGGTCCCATCTTTACAGG | Steric | MMMMMMMMMMMMMMMMMMMM |
| ASO17 | GAGGCACCAACTACAAAGAT | Gapmer | MMMMMDDDDDDDDDDMMMMM |
| ASO18 | GAGGCACCAACTACAAAGAT | Steric | MMMMMMMMMMMMMMMMMMMM |
| ASO19 | GACAGTGCTCCTAGACCAAA | Gapmer | MMMMMDDDDDDDDDDMMMMM |
| ASO20 | GACAGTGCTCCTAGACCAAA | Steric | MMMMMMMMMMMMMMMMMMMM |
| ASO21 | TAGTCACAGGGAGCATTAGG | Gapmer | MMMMMDDDDDDDDDDMMMMM |
| ASO22 | TAGTCACAGGGAGCATTAGG | Steric | MMMMMMMMMMMMMMMMMMMM |
| ASO23 | TAATGACCCAACCTTGTGTC | Gapmer | MMMMMDDDDDDDDDDMMMMM |
| ASO24 | TAATGACCCAACCTTGTGTC | Steric | MMMMMMMMMMMMMMMMMMMM |
| ASO25 | GATTAGGAAATGCACAACAC | Gapmer | MMMMMDDDDDDDDDDMMMMM |
| ASO26 | GATTAGGAAATGCACAACAC | Steric | MMMMMMMMMMMMMMMMMMMM |
| ASO27 | CAAGTTTCCATACCTGGTTC | Gapmer | MMMMMDDDDDDDDDDMMMMM |
| ASO28 | CAAGTTTCCATACCTGGTTC | Steric | MMMMMMMMMMMMMMMMMMMM |
| ASO29 | TTTTGAGCTTAGATATGGAC | Gapmer | MMMMMDDDDDDDDDDMMMMM |
| ASO30 | TTTTGAGCTTAGATATGGAC | Steric | MMMMMMMMMMMMMMMMMMMM |
| ASO31 | GATTTTAAGCAGAATCCAGA | Gapmer | MMMMMDDDDDDDDDDMMMMM |
| ASO32 | GATTTTAAGCAGAATCCAGA | Steric | MMMMMMMMMMMMMMMMMMMM |
| ASO33 | CAAGTATAATCTCGCTTCTC | Gapmer | MMMMMDDDDDDDDDDMMMMM |
| ASO34 | CAAGTATAATCTCGCTTCTC | Steric | MMMMMMMMMMMMMMMMMMMM |
| ASO35 | GGTAGACAGGGTCCTTCATA | Gapmer | MMMMMDDDDDDDDDDMMMMM |
| ASO36 | GGTAGACAGGGTCCTTCATA | Steric | MMMMMMMMMMMMMMMMMMMM |

| | | | |
|---|---|---|---|
| ASO37 | CTGCCATACCCTTTCAATTG | Gapmer | MMMMMDDDDDDDDDDMMMMM |
| ASO38 | CTGCCATACCCTTTCAATTG | Steric | MMMMMMMMMMMMMMMMMMMM |
| ASO39 | CTTGCTACATCACCACTAGG | Gapmer | MMMMMDDDDDDDDDDMMMMM |
| ASO40 | CTTGCTACATCACCACTAGG | Steric | MMMMMMMMMMMMMMMMMMMM |
| ASO41 | GGTAATGGTAGTTTGTCTAA | Gapmer | MMMMMDDDDDDDDDDMMMMM |
| ASO42 | GGTAATGGTAGTTTGTCTAA | Steric | MMMMMMMMMMMMMMMMMMMM |
| ASO43 | GCGATCTGAGAGTTACTTTC | Gapmer | MMMMMDDDDDDDDDDMMMMM |
| ASO44 | GCGATCTGAGAGTTACTTTC | Steric | MMMMMMMMMMMMMMMMMMMM |
| ASO45 | TTTTTTCTCTCCACGTGTGT | Gapmer | MMMMMDDDDDDDDDDMMMMM |
| ASO46 | TTTTTTCTCTCCACGTGTGT | Steric | MMMMMMMMMMMMMMMMMMMM |
| ASO47 | GTGTGGAAACTGGCAATAAG | Gapmer | MMMMMDDDDDDDDDDMMMMM |
| ASO48 | GTGTGGAAACTGGCAATAAG | Steric | MMMMMMMMMMMMMMMMMMMM |
| ASO49 | TATTGTTTTGCGGCTTGGAC | Gapmer | MMMMMDDDDDDDDDDMMMMM |
| ASO50 | TATTGTTTTGCGGCTTGGAC | Steric | MMMMMMMMMMMMMMMMMMMM |
| ASO51 | TCTAACGTGCTGAAGGACCC | Gapmer | MMMMMDDDDDDDDDDMMMMM |
| ASO52 | TCTAACGTGCTGAAGGACCC | Steric | MMMMMMMMMMMMMMMMMMMM |
| ASO53 | GTCTAAGGCTTGAGGTTGAG | Gapmer | MMMMMDDDDDDDDDDMMMMM |
| ASO54 | GTCTAAGGCTTGAGGTTGAG | Steric | MMMMMMMMMMMMMMMMMMMM |
| ASO55 | TTCAGATCTGTGATCCACTG | Gapmer | MMMMMDDDDDDDDDDMMMMM |
| ASO56 | TTCAGATCTGTGATCCACTG | Steric | MMMMMMMMMMMMMMMMMMMM |
| ASO57 | AAGATTCTCTCCCTATGTCT | Gapmer | MMMMMDDDDDDDDDDMMMMM |
| ASO58 | AAGATTCTCTCCCTATGTCT | Steric | MMMMMMMMMMMMMMMMMMMM |
| ASO59 | GTTGAACTCTTTGCATAACC | Gapmer | MMMMMDDDDDDDDDDMMMMM |
| ASO60 | GTTGAACTCTTTGCATAACC | Steric | MMMMMMMMMMMMMMMMMMMM |
| ASO61 | CTCCTGACTATGTTTTTCAC | Gapmer | MMMMMDDDDDDDDDDMMMMM |
| ASO62 | CTCCTGACTATGTTTTTCAC | Steric | MMMMMMMMMMMMMMMMMMMM |
| ASO63 | GTGTTTTCCCAGTCTGTTGC | Gapmer | MMMMMDDDDDDDDDDMMMMM |
| ASO64 | GTGTTTTCCCAGTCTGTTGC | Steric | MMMMMMMMMMMMMMMMMMMM |
| ASO65 | CGAGGTTCTCTTTCAAGGAT | Gapmer | MMMMMDDDDDDDDDDMMMMM |
| ASO66 | TGAGATGCAAAGCACGAGGT | Gapmer | MMMMMDDDDDDDDDDMMMMM |
| ASO67 | GTCAGAGGCAGTTGAGATGC | Gapmer | MMMMMDDDDDDDDDDMMMMM |
| ASO68 | ACCCAACCTTGTGTCAGAGG | Gapmer | MMMMMDDDDDDDDDDMMMMM |
| ASO69 | ATGACCCAACCTTGTGTCAG | Gapmer | MMMMMDDDDDDDDDDMMMMM |
| ASO70 | GGTTAATGACCCAACCTTGT | Gapmer | MMMMMDDDDDDDDDDMMMMM |
| ASO71 | ATATGGTTAATGACCCAACC | Gapmer | MMMMMDDDDDDDDDDMMMMM |
| ASO72 | TATGAAGTAAGAAAGGGGTA | Gapmer | MMMMMDDDDDDDDDDMMMMM |
| ASO73 | GACAAGATTAGGAAATGCAC | Gapmer | MMMMMDDDDDDDDDDMMMMM |
| ASO74 | CACAGAAGACAAGATTAGGA | Gapmer | MMMMMDDDDDDDDDDMMMMM |
| ASO75 | TCTGCACAGAAGACAAGATT | Gapmer | MMMMMDDDDDDDDDDMMMMM |

| | | | |
|---|---|---|---|
| ASO76 | TTAATACAGCTCTGGAGTGGGGT | Gapmer | MMMMMDDDDDDDDDDDDDMMMMM |
| ASO77 | TTAATACAGCTCTGGAGTGGGGT | Mixmer | MMMMMMDDDMDDDMDDDMMMMMM |
| ASO78 | TTAATACAGCTCTGGAGTGGGGT | Steric | MMMMMMMMMMMMMMMMMMMMMM M |
| ASO79 | TTAATACAGCTCTGGAGTGGGGT | Mixmer | MLMLMLDLDLDLDLDLDLMLMLM |
| ASO80 | TTAATACAGCTCTGGAGTGGGGT | Mixmer | MMLMMLMMLMMLMMLMMLMMLMM |
| ASO81 | TATGAAGTAAGAAAGGGGTA | Steric | MMMMMMMMMMMMMMMMMMMM |
| ASO82 | TATGAAGTAAGAAAGGGGTA | Mixmer | MMMLMDDLDDDLDDDLMMMM |
| ASO83 | TATGAAGTAAGAAAGGGGTA | Mixmer | MMMMMLMMMMLMMMMLMMMM |
| NTC ASO 1 | AAGGTTCGGAGTCGTAGCTG | Gapmer | MMMMMDDDDDDDDDDMMMMM |
| NTC ASO 2 | AAGGTTCGGAGTCGTAGCTG | Steric | MMMMMMMMMMMMMMMMMMMM |
| NTC ASO 3 | AAGGTTCGGAGTCGTAGCTG | Mixmer | MMMLMMMLMMMLMMMLMMMM |

M MOE
D DNA
L LNA
*All linkages are phosphorothioate unless otherwise noted

**Supplementary Table 4: Antibodies**

| Target | Vendor | Catalog number |
|---|---|---|
| ACTB | RD System | MAB8929 |
| ARID1A | Santa Cruz Biotechnology | sc-32761X |
| BCL6 | Life Technologies | PA527390 |
| H3K27ac | Abcam | ab4729 |
| H3K4me1 | Abcam | ab8895 |
| H3K4me3 | Abcam | ab8580 |
| HDAC1 | Active Motif | 40967ACTMOTIF |
| HDAC5 | Active Motif | 40970ACTMOTIF |
| NCOR | Life Technologies | A301145A |
| OTC | Abcam | ab203859 |
| SP1 | Santa Cruz Biotechnology | sc-17824X |
| Phalloidin-Alexa Fluor | Life Technologies | A22287 |
| VCL | Cell Signaling Technologies | 13901S |

**Supplementary Table 5: Custom primers/probes and sequences**

| **cdPCR primers/probes for regRNA Quantification** | | |
|---|---|---|
| **Primer name** | **Species** | **Sequence (5' → 3')** |
| *TERC* Forward | Hg38 | AAGAGGAACGGAGCGAGTC |
| *TERC* Reverse | Hg38 | CACCAACAGGAAAGCGAACT |
| *TERC* Probe | Hg38 | ATTCCCTGAGCTGTGGGACGTG |
| *OTC* regRNA + Strand For. | Hg38 | TTTGATGATTGGCATTTCACA |
| *OTC* regRNA + Strand Rev. | Hg38 | ATCCCTTTTCCTGCCATACC |
| *OTC* regRNA + Strand Probe | Hg38 | GCCTTTATGAAGGACCCTGTC |
| *OTC* regRNA - Strand For. | Hg38 | TGGATTTCCAGGTCACCTTC |
| *OTC* regRNA - Strand Rev. | Hg38 | GCTAGTACTGCTTGCCAGATGA |
| *OTC* regRNA - Strand Probe | Hg38 | TGCAATGTATTACCATTAAGAAGTAAGCGA |
| *OTC* regRNA + spliced with mRNA For. | Hg38 | CTCAACTGCCTCTGACACAA |
| *OTC* regRNA + spliced with mRNA Rev. | Hg38 | CAAGGGCATAGAATCGTCCTT |
| **ChIP-qPCR primers** | | |
| *OTC* enh summit For. | Hg38 | |
| *OTC* enh summit Rev. | Hg38 | |
| GAPDH positive control For. | Hg38 | |
| GAPDH positive control Rev. | Hg38 | |
| Negative control For. | Hg38 | |
| Negative control Rev. | Hg38 | |
| **Cloning sequences** | | |
| *OTC* minus strand regRNA (IVT) | Hg38 | CACAGAGCAAACAGGGAAATCACAGAGGTTCAAAGTTCACCA<br>GTGTCTCAACAATCAGCCCTAATGCTCCCTGTGACTACCAGAC<br>ACTCCCAGGACCTGAGTGATGGGGATCCCATGAGATCATTTTC<br>ATTGCTTTCTACTGACCAGGGTTTGGTCTAGGAGCACTGTCCC<br>AGTAATAATTTTCATGGCAATATTCTCCCCTTGAGCCCAGGAAA<br>CATGTCTTGGATGGCTTCAAAAGTCACTGCTTGGTGAATGCCT<br>TCTCTGCCCATTTCTACTTTTTGGTGAAACTTGAAACCATCTTT<br>GTAGTTGGTGCCTCTCTTCAGACCCTACTTGGGAGGTGCTCTT<br>GACCTGCTATTGATTGCTTTATTGGGCTATATCTACTAAGCAGG<br>GGCTCTGCCCTCACCTTAAGCTAATGATTAAACACAGCCTTCTT<br>CTCTCAAGGCTGCTCCACTGGTAACAACTCTGTGGCCTGTAAA<br>GATGGGACCTATTTAGGGTCTGGAAGATAGACCATGGGAATCC<br>TGTCTTCAAGATTCAAGAGAAACAAGCCCTTTTCATGGGGCTT<br>TGTTGAGTGTTTGGAGCCTAGGTCATAGGTGCTACATATTCACC<br>ATTATTGATTTATTCCTCCAGAATTTTTCAACTGGAGTTCACCAT<br>TTCTTCCAGGGAACCAAGGAGTTCATGGATTTCCAGGTCACCT<br>TCATTGTTATGCAATGTATTACCATTAAGAAGTAAGCGAATCATC<br>TGGCAAGCAGTACTAGCAGCTCCTACTCATAGCTTTGTTGTGA<br>GTATGAAATGTAATAATGAATAGAGAGTACTGTAGCACAGTACC<br>TAGCTCAGTGTTCAATAAATGTTAGCTTTTGTTAACTACTACCAT<br>TGGCACATGTGGTGAGAGGCCCCATCCCTGGCTCAGTTCTTG<br>GCTTATTCTAATCACTTTCCTACAAATAAAAGTGTTGAGGTGTC<br>CGTCTTTCTTTCATACCCCCACCCCACTCCAGAGCTGTATTAAA<br>AGTGAAATTCAGGCTGGGCATGGTGGCTCACGCCTGTAATCC |

| | | |
|---|---|---|
| | | CAGCACTTTGAGGCGGGCGGATCACGAGGTCAGGAGTTCGA<br>GACTAGCCTGACCAACGTGGTGAAACCCCGGCTCTACTAAAAA<br>TACAAAAATTAGCCAGGCATGGTGGCGGACACCTGTAATCCCA<br>GCTATGCATCGAGAGGCTGAGGCAGGAGAATTGCTTGAACCC<br>GGGAGGCGGAGGTTGCAGTGAGCCGAGATAGTGCCACTGCT<br>CTCTAGCCTGGGCGACAGAGCGAGACACCATCTCC |
| *OTC* minus strand regRNA (mutated/scrambled ASO1 position; IVT) | Hg38 | CACAGAGCAAACAGGGAAATCACAGAGGTTCAAAGTTCACCAGTGTCT<br>CAACAATCAGCCCTAATGCTCCCTGTGACTACCAGACACTCCCAGGAC<br>CTGAGTGATGGGGATCCCATGAGATCATTTTCATTGCTTTCTACTGACCA<br>GGGTTTGGTCTAGGAGCACTGTCCCAGTAATAATTTTCATGGCAATATTCT<br>CCCCTTGAGCCCAGGAAACATGTCTTGGATGGCTTCAAAAGTCACTGCT<br>TGGTGAATGCCTTCTCTGCCCATTTCTACTTTTTGGTGAAACTTGAAACCA<br>TCTTTGTAGTTGGTGCCTCTCTTCAGACCCTACTTGGGAGGTGCTCTTGA<br>CCTGCTATTGATTGCTTTATTGGGCTATATCTACTAAGCAGGGGCTCTGCC<br>CTCACCTTAAGCTAATGATTAAACACAGCCTTCTTCTCTCAAGGCTGCTC<br>CACTGGTAACAACTCTGTGGCCTGTAAAGATGGGACCTATTTAGGGTCTG<br>GAAGATAGACCATGGGAATCCTGTCTTCAAGATTCAAGAGAAACAAGCC<br>CTTTTCATGGGGCTTTGTTGAGTGTTTGGAGCCTAGGTCATAGGTGCTAC<br>ATATTCACCATTATTGATTTATTCCTCCAGAATTTTTCAACTGGAGTTCACC<br>ATTTCTTCCAGGGAACCAAGGAGTTCATGGATTTCCAGGTCACCTTCATT<br>GTTATGCAATGTATTACCATTAAGAAGTAAGCGAATCATCTGGCAAGCAGT<br>ACTAGCAGCTCCTACTCATAGCTTTGTTGTGAGTATGAAATGTAATAATGAA<br>TAGAGAGTACTGTAGCACAGTACCTAGCTCAGTGTTCAATAAATGTTAGCT<br>TTTGTTAACTACTACCATTGGCACATGTGGTGAGAGGCCCCATCCCTGG<br>CTCAGTTCTTGGCTTATTCTAATCACTTTCCTACAAATAAAAGTGTTGAGGT<br>GTCCGTCTTTCTTTCATACCCCCACCTTGTCTTGAGATCACGCCGGAAGT<br>GAAATTCAGGCTGGGCATGGTGGCTCACGCCTGTAATCCCAGCACTTT<br>GAGGCGGGCGGATCACGAGGTCAGGAGTTCGAGACTAGCCTGACCAA<br>CGTGGTGAAACCCCGGCTCTACTAAAAATACAAAAATTAGCCAGGCATG<br>GTGGCGGACACCTGTAATCCCAGCTATGCATCGAGAGGCTGAGGCAG<br>GAGAATTGCTTGAACCCGGGAGGCGGAGGTTGCAGTGAGCCGAGATA<br>GTGCCACTGCTCTCTAGCCTGGGCGACAGAGCGAGACACCATCTCC |
| *OTC* enhancer (region 2) reporter assay | Hg38 | TTATTTTTTATGTTTTTTGTTTGTTTGTTTGTTTGAGACAGAGTC<br>TCGCTCTGTCACCCAGGCTGGAATGCAGTGGCACAACCTCGG<br>CTCATGGCAACCTCTGCCTCCTGGGTTCAAGCGATCTGAGAG<br>TTACTTTCTAATTAACAGATGGTAATGGTAATGGTAGTTTGTCTA<br>AAAATATTGATTCAGGGGTGTTTTCTTGCTACATCACCACTAGG<br>TATTTATTCAAACAGGTGGGCCCTAGGGGAAATTGTGAAATCA<br>GTGAGGATTATCTCTGACCCCCTCTCCATCCCTTTTCCTGCCAT<br>ACCCTTTCAATTGAGGCATTTGAAGAAGGTAGACAGGGTCCTT<br>CATAAAGGCTAAAATTCTGAACAGAGTGTGAAATGCCAATCATC<br>AAATTCAAGTATAATCTCGCTTCTCTCTTGTCATATAGACCACAT<br>TTCCATTGAAATCATTATTCTGTTAATCATTTGCCCTGGAAGTTA<br>TTGATTTTAAGCAGAATCCAGAATTTTTTGAGCTTAGATATGGAC<br>AAAAGTCTCAAGTTTCCATACCTGGTTCTTTTTATTTTGTGATCT<br>GGTAAATTTCTGCACAGAAGACAAGATTAGGAAATGCACAACA<br>CAATTTATGAAGTAAGAAAGGGGTAAATAAATATGGTTAATGACC<br>CAACCTTGTGTCAGAGGCAGTTGAGATGCAAAGCACGAGGTT<br>CTCTTTCAAGGATGCCCTTTTATCTCTTCAAGTTCCCACACAGA<br>ATCCTAGAGCTATCTCTTCTCCCATAGGCTGATTCCAATTCCTC<br>CCCATTGGAAGTACAGAGGTTAAACTGCATAAAGGAGATTTGTT<br>AGCAGACTGAAAGCTATCAGGCACAGAGCAAACAGGGAAATC<br>ACAGAGGTTCAAAGTTCACCAGTGTCTCAACAATCAGCCCTAA<br>TGCTCCCTGTGACTACCAGACACTCCCAGGACCTGAGTGATG<br>GGGATCCCATGAGATCATTTTCATTGCTTTCTACTGACCAGGGT<br>TTGGTCTAGGAGCACTGTCCCAGTAATAATTTTCATGGCAATAT<br>TCTCCCCTTGAGCCCAGGAAACATGTCTTGGATGGCTTCAAAA<br>GTCACTGCTTGGTGAATGCCTTCTCTGCCCATTTCTACTTTTTG<br>GTGAAACTTGAAACCATCTTTGTAGTTGGTGCCTCTCTTCAGA<br>CCCTACTTGGGAGGTGCTCTTGACCTGCTATTGATTGCTTTATT |

| | | |
|---|---|---|
| | | GGGCTATATCTACTAAGCAGGGGCTCTGCCCTCACCTTAAGCT<br>AATGATTAAACACAGCCTTCTTCTCTCAAGGCTGCTCCACTGG<br>TAACAACTCTGTGGCCTGTAAAGATGGGACCTATTTAGGGTCT<br>GGAAGATAGACCATGGGAATCCTGTCTTCAAGATTCAAGAGAA<br>ACAAGCCCTTTTCATGGGGCTTTGTTGAGTGTTTGGAGCCTAG<br>GTCATAGGTGCTACATATTCACCATTATTGATTTATTCCTCCAGA<br>ATTTTTCAACTGGAGTTCACCATTTCTTCCAGGGAACCAAGGA<br>GTTCATGGATTTCCAGGTCACCTTCATTGTTATGCAATGTATTAC<br>CATTAAGAAGTAAGCGAATCATCTGGCAAGCAGTACTAGCAGC<br>TCCTACTCATAGCTTTGTTGTGAGTATGAAATGTAATAATGAATA<br>GAGAGTACTGTAGCACAGTACCTAGCTCAGTGTTCAATAAATGT<br>TAGCTTTTGTTAACTACTACCATTGGCACATGTGGTGAGAGGC<br>CCCATCCCTGGCTCAGTTCTTGGCTTATTCTAATCACTTTCCTA<br>CAAATAAAAGTGTTGAGGTGTCCGTCTTTCTTTCATACCCCCAC<br>CCCACTCCAGAGCTGTATTAAA |
| *OTC* enhancer (region 3) reporter assay | Hg38 | TATTTTTTATGTTTTTTGTTTGTTTGTTTGTTTGAGACAGAGTCT<br>CGCTCTGTCACCCAGGCTGGAATGCAGTGGCACAACCTCGGC<br>TCATGGCAACCTCTGCCTCCTGGGTTCAAGCGATCTGAGAGTT<br>ACTTTCTAATTAACAGATGGTAATGGTAATGGTAGTTTGTCTAAA<br>AATATTGATTCAGGGGTGTTTTCTTGCTACATCACCACTAGGTA<br>TTTATTCAAACAGGTGGGCCCTAGGGGAAATTGTGAAATCAGT<br>GAGGATTATCTCTGACCCCCTCTCCATCCCTTTTCCTGCCATAC<br>CCTTTCAATTGAGGCATTTGAAGAAGGTAGACAGGGTCCTTCA<br>TAAAGGCTAAAATTCTGAACAGAGTGTGAAATGCCAATCATCAA<br>ATTCAAGTATAATCTCGCTTCTCTCTTGTCATATAGACCACATTT<br>CCATTGAAATCATTATTCTGTTAATCATTTGCCCTGGAAGTTATT<br>GATTTTAAGCAGAATCCAGAATTTTTTGAGCTTAGATATGGACA<br>AAAGTCTCAAGTTTCCATACCTGGTTCTTTTTATTTTGTGATCTG<br>GTAAATTTCTGCACAGAAGACAAGATTAGGAAATGCACAACAC<br>AATTTATGAAGTAAGAAAGGGGTAAATAAATATGGTTAATGACCC<br>AACCTTGTGTCAGAGGCAGTTGAGATGCAAAGCACGAGGTTC<br>TCTTTCAAGGATGCCCTTTTATCTCTTCAAGTTCCCACACAGAA<br>TCCTAGAGCTATCTCTTCTCCCATAGGCTGATTCCAATTCCTCC<br>CCATTGGAAGTACAGAGGTTAAACTGCATAAAGGAGATTTGTTA<br>GCAGACTGAAAGCTATCAGGCACAGAGCAAACAGGGAAATCA<br>CAGAGGTTCAAAGTTCACCAGTGTCTCAACAATCAGCCCTAAT<br>GCTCCCTGTGACTACCAGACACTCCCAGGACCTGAGTGATGG<br>GGATCCCATGAGATCATTTTCATTGCTTTCTACTGACCAGGGTT<br>TGGTCTAGGAGCACTGTCCCAGTAATAATTTTCATGGCAATATT<br>CTCCCCTTGAGCCCAGGAAACATGTCTTGGATGGCTTCAAAAG<br>TCACTGCTTGGTGAATGCCTTCTCTGCCCATTTCTACTTTTTGG<br>TGAAACTTGAAACCATCTTTGTAGTTGGTGCCTCTCTTCAGAC<br>CCTACTTGGGAGGTGCTCTTGACCTGCTATTGATTGCTTTATTG<br>GGCTATATCTACTAAGCAGGGGCTCTGCCCTCACCTTAAGCTA<br>ATGATTAAACACAGCCTTCTTCTCTCAAGGCTGCTCCACTGGT<br>AACAACTCTGTGGCCTGTAAAGATGGGACCTATTTAGGGTCTG<br>GAAGATAGACCATGGGAATCCTGTCTTCAAGATTCAAGAGAAA<br>CAAGCCCTTTTCATGGGGCTTTGTTGAGTGTTTGGAGCCTAGG<br>TCATAGGTGCTACATATTCACCATTATTGATTTATTCCTCCAGAA<br>TTTTTCAACTGGAGTTCACCATTTCTTCCAGGGAACCAAGGAG<br>TTCATGGATTTCCAGGTCACCTTCATTGTTATGCAATGTATT |
| *OTC* enhancer (region 4) reporter assay | Hg38 | TATTTTTTATGTTTTTTGTTTGTTTGTTTGTTTGAGACAGAGTCT<br>CGCTCTGTCACCCAGGCTGGAATGCAGTGGCACAACCTCGGC<br>TCATGGCAACCTCTGCCTCCTGGGTTCAAGCGATCTGAGAGTT<br>ACTTTCTAATTAACAGATGGTAATGGTAATGGTAGTTTGTCTAAA<br>AATATTGATTCAGGGGTGTTTTCTTGCTACATCACCACTAGGTA<br>TTTATTCAAACAGGTGGGCCCTAGGGGAAATTGTGAAATCAGT<br>GAGGATTATCTCTGACCCCCTCTCCATCCCTTTTCCTGCCATAC<br>CCTTTCAATTGAGGCATTTGAAGAAGGTAGACAGGGTCCTTCA<br>TAAAGGCTAAAATTCTGAACAGAGTGTGAAATGCCAATCATCAA<br>ATTCAAGTATAATCTCGCTTCTCTCTTGTCATATAGACCACATTT<br>CCATTGAAATCATTATTCTGTTAATCATTTGCCCTGGAAGTTATT |

| | | |
|---|---|---|
| | | GATTTTAAGCAGAATCCAGAATTTTTTGAGCTTAGATATGGACA<br>AAAGTCTCAAGTTTCCATACCTGGTTCTTTTTATTTTGTGATCTG<br>GTAAATTTCTGCACAGAAGACAAGATTAGGAAATGCACAACAC<br>AATTTATGAAGTAAGAAAGGGGTAAATAAATATGGTTAATGACCC<br>AACCTTGTGTCAGAGGCAGTTGAGATGCAAAGCACGAGGTTC<br>TCTTTCAAGGATGCCCTTTTATCTCTTCAAGTTCCCACACAGAA<br>TCCTAGAGCTATCTCTTCTCCCATAGGCTGATTCCAATTCCTCC<br>CCATTGGAAGTACAGAGGTTAAACTGCATAAAGGAGATTTGTTA<br>GCAGACTGAAAGCTATCAGGCACAGAGCAAACAGGGAAATCA<br>CAGAGGTTCAAAGTTCACCAGTGTCTCAACAATCAGCCCTAAT<br>GCTCCCTGTGACTACCAGACACTCCCAGGACCTGAGTGATGG<br>GGATCCCATGAGATCATTTTCATTGCTTTCTACTGACCAGGGTT<br>TGGTCTAGGAGCACTGTCCCAGTAATAATTTTCATGGCAATATT<br>CTCCCCTTGAGCCCAGGAAACATGTCTTGGATGGCTTCAAAAG<br>TCACTGCTTGGTGAATGCCTTCTCTGCCCATTTCTACTTTTTGG<br>TGAAACTTGAAACCATCTTTGTAGTTGGTGCCTCTCTTCAGAC<br>CCTACTTGGGAGGTGCTCTTGACCTGCTATTGATTGCTTTATTG<br>GGCTATATCTACTAAGCAGGGGCTCTGCCCTCACCTTAAGCTA<br>ATGATTAAACACAGC |

**Supplementary Table 6: siRNA sequences**

| Target | siRNA sequence(s) | Manufacturer | Catalog number |
|---|---|---|---|
| *RNaseH1* | GCGCAGAGCCGUAUGCAAA<br>GAGCUAAACAAUCGGAAGA<br>GCCAGGCCAUCCUUUAAAU<br>GACAUUCAGUGGAUGCAUG | Dharmacon | siGENOME Human RNASEH1 (246243) siRNA - SMARTpool, 5nmol (M-012595-00-0005) |
| *HDAC1* | ACUAUGGUCUCUACCGAAA<br>GCAAGUAUUAUGCUGUUAA<br>CCGGUCAUGUCCAAAGUAA<br>CCACAGCGAUGACUACAUU | Dharmacon | ON-TARGETplus Human HDAC1 (3065) siRNA - SMARTpool, 10nmol (L-003493-00-0010) |
| *HDAC3* | GGAAAGCGAUGUGGAGAUU<br>AAAGCGAUGUGGAGAUUUA<br>GCAUUGAUGACCAGAGUUA<br>GGAAUGCGUUGAAUAUGUC | Dharmacon | siGENOME Human HDAC3 (8841) siRNA - SMARTpool, 5nmol (M-003496-02-0005) |
| *HDAC5* | GUUAUUAGCACCUUUAAGA<br>AAAGUGCGUUCAAGGCUAA<br>CAGCAGAGCACCCUCAUUG<br>GAAUUCCUCUUGUCGAAGU | Dharmacon | siGENOME Human HDAC5 (10014) siRNA - SMARTpool, 5 nmol (M-003498-02-0) |
| *NCOR1* | GAUCACAUCUGUCAAAUUA<br>GAACGUGGCUCUCAAAGUU<br>GAAAGGAAAUCGACACUGA<br>GCCCUGGGAUUUAUGAUGA | Dharmacon | siGENOME Human NCOR1(9611) siRNA - SMARTpool, 5 nmol (SO-3030898G) |
| *NCOR2* | GGACGGAGAUCUUCAAUAU<br>GAACCUCGAUGAGAUCUUG<br>GGAAAAGACUCAAAGUAAA<br>GCGCACCUAUGACAUGAUG | Dharmacon | siGENOME Human NCOR2 (9612) siRNA - SMARTpool, 5nmol (M-020145-02-0005) |
| Non-targeting | UAGCGACUAAACACAUCAA<br>UAAGGCUAUGAAGAGAUAC<br>AUGUAUUGGCCUGUAUUAG<br>AUGAACGUGAAUUGCUCAA | Dharmacon | siGENOME Non-Targeting siRNA Pool #1, 5nmol (D-001206-13-05) |
| Non-targeting | UGGUUUACAUGUCGACUAA<br>UGGUUUACAUGUUGUGUGA<br>UGGUUUACAUGUUUUCUGA<br>UGGUUUACAUGUUUUCCUA | Dharmacon | ON-TARGETplus Non-targeting Pool, 5 nmol (D-001810-10-05) |

| Dataset | Replicate | Treatment | Source | Type | Identifier |
|---|---|---|---|---|---|
| H3K27ac ChIP-seq (PHH) | Rep1 | -- | This study | ChIP-seq | |
| | Rep2 | -- | This study | ChIP-seq | |
| H3K4me3 ChIP-seq (PHH) | -- | -- | This study | ChIP-seq | |
| H3K4me1 ChIP-seq (PHH) | -- | -- | This study | ChIP-seq | |
| ATAC-seq (PHH) | Rep1 | -- | This study | ATAC-seq | |
| | Rep2 | -- | This study | ATAC-seq | |
| PRO-seq (PHH) | Rep1 | -- | This study | PRO-seq | |
| | Rep2 | -- | This study | PRO-seq | |
| regRNA capture-seq (VeryHigh) | PolyA | -- | This study | PacBio seq | |
| | Non-polyA | -- | This study | PacBio seq | |
| regRNA capture-seq (High) | PolyA | -- | This study | PacBio seq | |
| | Non-polyA | -- | This study | PacBio seq | |
| regRNA capture-seq (Medium) | PolyA | -- | This study | PacBio seq | |
| | Non-polyA | -- | This study | PacBio seq | |
| regRNA capture-seq (Low) | PolyA | -- | This study | PacBio seq | |
| | Non-polyA | -- | This study | PacBio seq | |
| regRNA capture-seq (*OTC*, PHH) | PolyA | -- | This study | PacBio seq | |
| | Non-polyA | -- | This study | PacBio seq | |
| PolyA regRNA capture-seq (*OTC*, liver) | PolyA | -- | This study | PacBio seq | |
| | Non-polyA | -- | This study | PacBio seq | |
| H3K27ac ChIP-seq (Liver) | -- | -- | GSE17312 | ChIP-seq | GSM1112809 |
| H3K4me3 ChIP-seq (Liver) | -- | -- | GSE16256 | ChIP-seq | |
| ATAC-seq (Liver) | -- | -- | ENCODE, ENCSR124NNL | ATAC-seq | |
| PolyA RNA-seq (PHH) | -- | -- | This study | RNA-seq | |
| rRNA-depleted RNA-seq (PHH) | -- | -- | This study | RNA-seq | |
| PRO-seq JAKi (PHH) | Rep1 | 3μM upadacitinib | This study | PRO-seq | |
| | Rep2 | 3μM upadacitinib | This study | PRO-seq | |
| PRO-seq DMSO (PHH) | Rep1 | DMSO | This study | PRO-seq | |
| | Rep2 | DMSO | This study | PRO-seq | |
| H3K4me3 ChIP-seq (HepG2) | -- | -- | ENCODE, ENCFF645ZUY | ChIP-seq | |
| H3K27ac ChIP-seq (HepG2) | -- | -- | ENCODE, ENCFF493VUL | ChIP-seq | |
| H3K4me1 ChIP-seq (HepG2) | -- | -- | ENCODE, ENCFF470RYT | ChIP-seq | |
| ATAC-seq (HepG2) | -- | -- | ENCODE, ENCFF285FQS | ATAC-seq | |
| SP1 ChIP-seq (HepG2) | -- | -- | ENCODE , ENCFF239YXQ | ChIP-seq | |
| BCL6 ChIP-seq (HepG2) | -- | -- | ENCODE, ENCFF648HUD | ChIP-seq | |
| NCOR1 ChIP-seq (HepG2) | -- | -- | ENCODE, ENCFF601PYD | ChIP-seq | |
| HDAC1 ChIP-seq (HepG2) | -- | -- | ENCODE, ENCFF180PDN | ChIP-seq | |
| ARID1A ChIP-seq (HepG2) | -- | -- | GSE69566 | ChIP-seq | |
| OTC regRNA SHAPE-MaP (+1M7) | -- | -- | This study | SHAPE-Map | |
| OTC regRNA SHAPE-MaP (+DMSO) | -- | -- | This study | SHAPE-Map | |
| OTC regRNA SHAPE-MaP (denatured) | -- | -- | This study | SHAPE-Map | |

| | | | | |
|---|---|---|---|---|
| OTC regRNA SHAPE-MaP + (+1M7) | -- | ASO1 | This study | SHAPE-Map |
| OTC regRNA SHAPE-MaP (+DMSO) | -- | ASO1 | This study | SHAPE-Map |
| OTC regRNA SHAPE-MaP (denatured) | -- | ASO1 | This study | SHAPE-Map |
| OTC regRNA SHAPE-MaP (+1M7) | -- | NTC ASO | This study | SHAPE-Map |
| OTC regRNA SHAPE-MaP (+DMSO) | -- | NTC ASO | This study | SHAPE-Map |
| OTC regRNA SHAPE-MaP (denatured) | -- | NTC ASO | This study | SHAPE-Map |
| Mutated OTC regRNA SHAPE-MaP (+1M7) | -- | ASO1 | This study | SHAPE-Map |
| Mutated OTC regRNA SHAPE-MaP (+DMSO) | -- | ASO1 | This study | SHAPE-Map |
| Mutated OTC regRNA SHAPE-MaP (denatured) | -- | ASO1 | This study | SHAPE-Map |
| Mutated OTC regRNA SHAPE-MaP (+1M7) | -- | NTC ASO | This study | SHAPE-Map |
| Mutated OTC regRNA SHAPE-MaP (+DMSO) | -- | NTC ASO | This study | SHAPE-Map |
| Mutated OTC regRNA SHAPE-MaP (denatured) | -- | NTC ASO | This study | SHAPE-Map |

*PHH, primary human hepatocyte;*
